# Simultaneous regulation of cytokinetic furrow and nucleus positions by cortical tension contributes to proper DNA segregation during late mitosis

**DOI:** 10.1101/626903

**Authors:** Anne Pacquelet, Matthieu Jousseaume, Jocelyn Etienne, Grégoire Michaux

**Affiliations:** Univ Rennes, CNRS, IGDR (Institut de Génétique et de Développement de Rennes), UMR 6290, F-35000 Rennes, France; Univ Grenoble Alpes, CNRS, LIPhy, 38000 Grenoble, France

## Abstract

Coordinating mitotic spindle and cytokinetic furrow positioning is essential to ensure proper DNA segregation. Here we present a novel mechanism, which corrects DNA segregation defects due to cytokinetic furrow mispositioning. We show that DNA segregation defects following the abnormal displacement of the cytokinetic furrow towards the anterior side of *C. elegans* one-cell embryos are unexpectedly corrected at the end of cytokinesis. This correction relies on the concomitant displacement of the furrow and of the anterior nucleus towards the posterior and anterior poles, respectively. It also coincides with cortical blebbing and an anteriorly directed flow of cytoplasmic particles. While microtubules contribute to nuclear displacement, relaxation of an excessive tension at the anterior cortex plays a central role in the correction process and simultaneously regulates cytoplasmic flow as well as nuclear and furrow displacements. This work thus reveals the existence of a so far uncharacterized correction mechanism, which is critical to correct DNA segregation defects due to cytokinetic furrow mispositioning.

## Introduction

Ensuring equal DNA segregation is a critical feature of cell division. The correct assembly of the mitotic spindle at metaphase followed by the separation of sister chromatids at anaphase are essential steps in this process. Ingression of the cytokinetic furrow between the separated chromatids then ensures that the genetic material is equally inherited by the two daughter cells. It is therefore crucial that the positions of the cytokinetic furrow and the mitotic spindle are well coordinated. Two main signalling pathways emanating from the mitotic spindle control the position of furrow ingression by restricting the localization of myosin contractile ring components to the vicinity of the central spindle (Bringmann and Hyman, 2005; Dechant and Glotzer, 2003): on the one hand, the centralspindlin complex, which localizes both at the central spindle and at the adjacent equatorial cortex, activates myosin contractility through the regulation of the small GTPase Rho (Basant et al., 2015; Nishimura and Yonemura, 2006; Yüce et al., 2005); on the other hand, astral microtubules appear to prevent myosin activity at the pole of dividing cells, through mechanisms that remain to be precisely determined (Bringmann et al., 2007; Lewellyn et al., 2010; Mangal et al., 2018; Price and Rose, 2017; Werner et al., 2007). Furthermore, several studies have shown that asymmetrically localized myosin also contributes to the localization of the site of furrow ingression (Cabernard et al., 2010; Ou et al., 2010; Pacquelet et al., 2015). Mistakes occurring at any of the successive steps of mitosis may impair DNA segregation. However, several control mechanisms limit the occurrence of detrimental DNA segregation defects. One of those mechanisms is the well characterized spindle assembly checkpoint, which inhibits entry into anaphase until all chromosomes are stably attached to the spindle (reviewed in Lara-Gonzalez et al., 2012). Similarly, at the end of mitosis, the abscission checkpoint delays abscission, for instance when chromosomes are lagging within the intracellular bridge (Norden et al., 2006; Steigemann et al., 2009). Furthermore, it has been shown in *Drosophila* neuroblasts that trailing chromatids can regulate cortical myosin to induce cell shape changes and ensure proper chromatid segregation (Kotadia et al., 2012; Montembault et al., 2017). DNA segregation defects may also result from the lack of coordination between the positions of the mitotic spindle and the cytokinetic furrow. Whether such DNA segregation defects can be corrected is so far unknown.

We have previously shown that preventing excessive cortical myosin activity is essential to properly localize the cytokinetic furrow during the first division of the one-cell *C. elegans* embryo (Pacquelet et al., 2015). During this division, myosin accumulates at the anterior cortex at early anaphase. This accumulation is transient and disappears later during anaphase while myosin localizes at the equatorial cortex, at the presumptive furrow ingression site. This dynamic localization of myosin needs to be tightly regulated. In particular, we found that the kinase PIG-1 and the anillin ANI-1 are required to restrict myosin anterior accumulation: in embryos lacking both PIG-1 and ANI-1 the accumulation of myosin at the anterior cortex is stronger and lasts longer than in control embryos. We have further shown that this abnormal accumulation of myosin leads to the displacement of the cytokinetic furrow towards the anterior of the embryo; as a result, the position of the cytokinetic furrow is no longer coordinated with the position of the mitotic spindle and does not initiate cleavage between the two sets of sister chromatids (Pacquelet et al., 2015).

In this study, we identify and characterize an unexpected rescue mechanism, which prevents DNA segregation defects resulting from the lack of coordination between cytokinetic furrow and mitotic spindle positions. Indeed, while furrow mispositioning first leads to DNA segregation defects in *ani-1;pig-1* embryos, we find that these defects are corrected during late mitosis. This correction involves both nuclear and furrow displacement. We find that microtubules/nucleus interactions contribute to nuclear displacement while myosin and cortical tension play a central role in the correction process by giving rise to cortical blebbing and cytoplasmic flow, which in turn drives both furrow and nuclear displacement.

## Results

### DNA segregation defects due to furrow mispositioning can be corrected

In *ani-1(RNAi);pig-1(gm344)* embryos (further referred to as *ani-1;pig-1)*, excessive myosin accumulation at the anterior cortex leads to the displacement of the furrow towards the anterior of the embryo (Fig.1A and movie 1) (Pacquelet et al., 2015). As a result, in most *ani-1;pig-1* embryos, the two nuclei are both located on the posterior side of the furrow, leading to strong DNA segregation defects (17/20 of *ani-1;pig-1* embryos recorded at 23°C). However, we surprisingly observed that most of those DNA segregation defects are corrected, with the anterior nucleus finally being located on the anterior side of the furrow (16/17 of *ani-1;pig-1* embryos with initial DNA segregation defects; Fig.1A and movie 1). In some of those embryos, correction of the DNA segregation defects is followed by quick furrow regression (3/16) but furrow ingression completes in 13/16 embryos and persists until the beginning of the second division in 8/16 embryos. As we observed that cytokinesis is delayed in *ani-1;pig-1* embryos (e.g. Fig.1A), we next asked when the correction of DNA segregation defects occurs relative to other cell cycle events. To this end, we first used the inner nuclear envelope protein EMR-1::mCherry to monitor nuclear envelope reformation. Both in control and *ani-1;pig-1* embryos, nuclear envelope already reassembles during cytokinesis (Fig.1B, movie 2). In *ani-1;pig-1* embryos, this always precedes the correction of DNA segregation defects (n=18/18). We next examined the mitotic spindle by using strains where the mitotic and the central spindle were labelled with a-tubulin::YFP or SPD-1::GFP, respectively. While spindle disassembly coincides with the end of furrow ingression in control embryos, it precedes the correction of DNA segregation defects in *ani-1;pig-1* embryos (movie 3) (n=11/12). During anaphase, both control and *ani-1;pig-1* embryos assemble a compact central spindle labelled with SPD-1::GFP (Fig.1C, movie 4). In control embryos, the central spindle remnant then associates with the ingressing furrow to form the midbody (Fig.1C, movie 4). In *ani-1;pig-1* embryos, it however disassembles before closure of the cytokinetic furrow and correction of DNA segregation defects (n=21/22) (Fig.1C, movie 4). Thus, the correction of DNA segregation defects that we observe in *ani-1;pig-1* embryos occur late during the cell cycle, when the nuclear envelope has already reformed and the mitotic and central spindle have disassembled.

**Figure 1:**
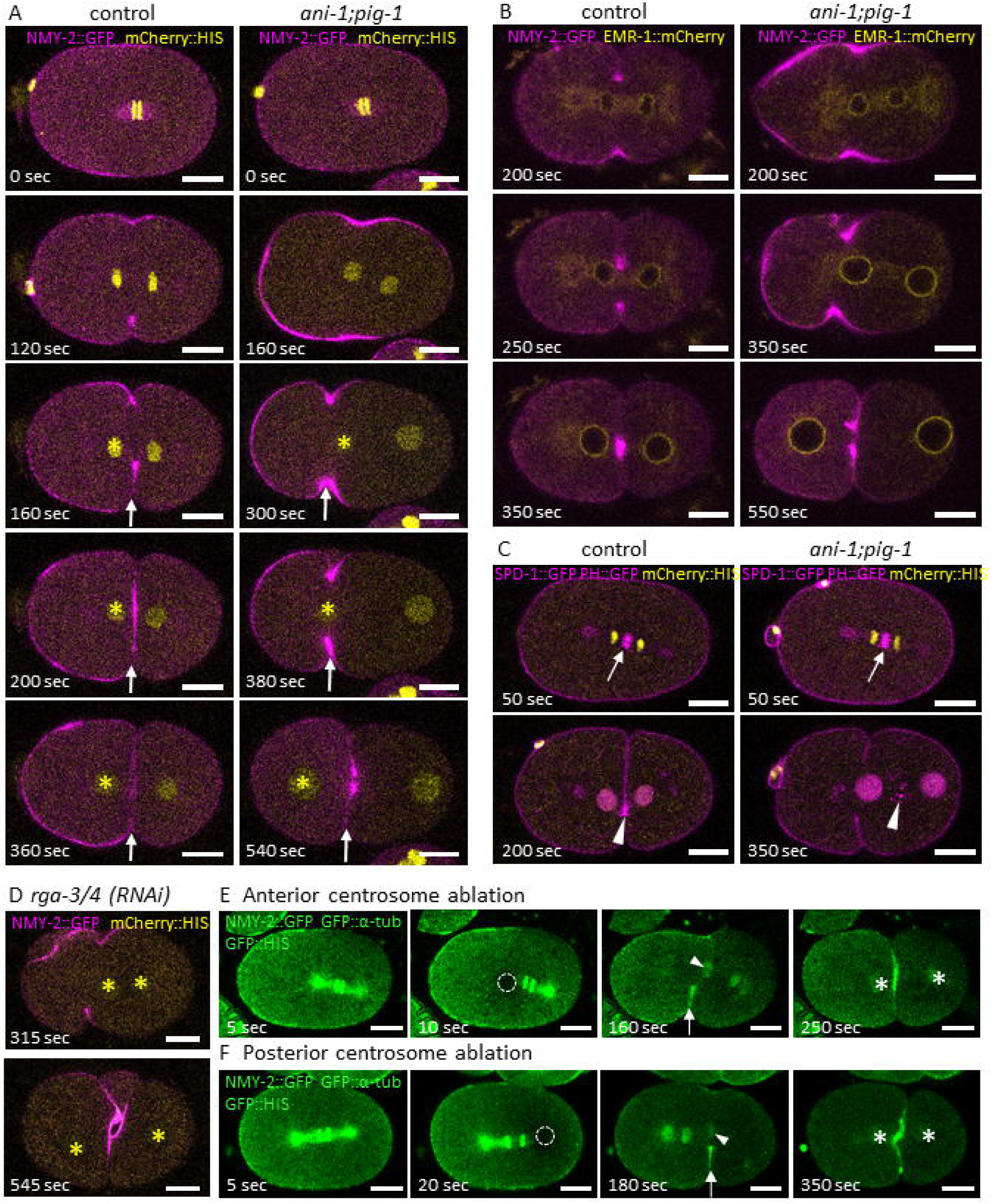
DNA segregation defects resulting from furrow mispositioning are corrected during late mitosis. **A.** Control and *ani-1(RNAi);pig-1(gm344)* embryos expressing NMY-2::GFP (magenta, labels non-muscle-myosin 2) and mCherry::HIS (yellow, labels DNA). Increased myosin levels at the anterior cortex of *ani-1(RNAi);pig-1(gm344)* embryos lead to furrow mispositioning and DNA segregation defects (e.g. t = 300 or 380 sec). Those defects are corrected late during mitosis (t = 540 sec). In control embryos the anterior nucleus (yellow asterix) is slightly displaced during cytokinesis and furrow position remains stable (arrow). Correction in *ani-1(RNAi);pig-1(gm344)* embryos involves the strong displacement of the anterior nucleus (yellow asterix) to the anterior and the displacement of the furrow (arrow) to the posterior. **B.** Control and *ani-1(RNAi);pig-1(gm344)* embryos expressing NMY-2::GFP (magenta) and EMR-1::mCherry (yellow, labels nuclear envelope). Nuclear envelope reassembles before the correction of DNA segregation defects. **C.** Control and *ani-1(RNAi);pig-1(gm344)* embryos expressing SPD-1::GFP (magenta, labels the central spindle, centrosomes and the nuclei after division (Nahaboo et al., 2015)), PH::GFP (magenta, membrane labelling) and mCherry::HIS (yellow). A compact central spindle forms at anaphase (arrow); at the end of cytokinesis, it associates with the cytokinetic furrow and forms the midbody in control embryos (arrowhead). In *ani-1;pig-1* embryos, the central spindle disassembles (arrowhead) before the correction of DNA segregation defects. **D.** *rga-3/4(RNAi)* leads to increased myosin activity, displacement of the furrow towards the anterior of the embryo and DNA segregation defects (t = 315 sec), which are then corrected (t = 545 sec). Embryo expresses NMY-2::GFP (magenta) and mCherry::HIS (yellow, DNA also indicated by yellow asterix). **E-F.** Anterior (E) and posterior (F) centrosome ablation lead to spindle and/or furrow mispositioning and DNA segregation defects in 15/16 and 18/33 embryos, respectively. These defects are then corrected in 14/15 and 17/18 embryos. Embryos express NMY-2::GFP (green, cortical and furrow signals), GFP::α-tubulin (green, labels the centrosome and to a lesser extent microtubules) and GFP::HIS (green, labels the DNA). Dashed circles indicate the ablated centrosomes. Arrowheads point to anterior (E) or posterior (F) DNA and arrows to the cytokinetic furrow before the correction of DNA segregation defects. Asterix indicate the position of the nuclei after division. In A and C-F: t_0_ = anaphase onset. In B: t_0_ = nuclear envelope breakdown. In all panels: embryos are oriented with the anterior to the left. Scale bar: 10 μm.

Interestingly, this correction mechanism is not specific to *ani-1;pig-1* embryos. Indeed, increased myosin activity following depletion of the Rho GAPs RGA-3/4 similarly leads to furrow displacement towards the anterior and to DNA segregation defects which are then corrected (10/12 embryos, Fig.1D, movie 5). Moreover, we induced DNA segregation defects by performing laser ablation of anaphase centrosomes: following ablation, the mitotic spindle is rapidly displaced in the direction opposite to the ablated centrosome; additionally, the cytokinetic furrow sometimes moves towards the ablated centrosome. This relative mispositioning of the mitotic spindle and cytokinetic furrow leads to DNA segregation defects which are then corrected (Fig.1E-F and legend for quantifications, movie 6). Notably, although DNA segregation defects resulting from anterior and posterior centrosome ablations occur in the opposite direction, both can be corrected. Altogether, those results show that DNA segregation defects due to furrow and/or spindle mispositioning can be corrected in different contexts, thereby revealing the existence of an unknown correction mechanism.

### Correction of DNA segregation defects involves both nuclear and furrow displacement

In a first attempt to understand the mechanisms underlying this correction process, we tracked the positions of both the anterior and posterior nuclei as well as the position of the cytokinetic furrow. In control embryos, both the anterior and posterior nuclei are displaced towards the poles of the embryo during and after cytokinesis while the position of the cytokinetic furrow remains stable during ring closure (Fig.1A and 2A-C, movie 1). In *ani-1;pig-1* embryos, the anterior and posterior nuclei also move towards the poles of the embryo. However, the anterior nucleus moves faster and further than in control embryos (Fig.1A and 2A, movie 1). This increased nuclear displacement is specific to the anterior nucleus as the kinetics of the posterior nucleus displacement is very similar in *ani-1;pig-1* and control embryos (Fig.2B). Furthermore, while the cytokinetic furrow initially moves towards the anterior in *ani-1;pig-1* embryos, it is then displaced towards the posterior during the correction of DNA segregation defects (Fig.1A and 2C-D, movie 1). Thus, the correction of DNA segregation defects due to furrow mispositioning in *ani-1;pig-1* embryos involves both the displacement of the anterior nucleus and the cytokinetic furrow towards the anterior and posterior pole of the embryo, respectively. Notably, these two displacements occur concomitantly (Fig.2D). Importantly, correction of DNA segregation defects following anterior centrosome ablations involves similar furrow and anterior nucleus displacements (Fig.2E,G, and movie 6); on the other hand, correction of DNA segregation defects due to posterior centrosome ablation is associated with displacements in the opposite direction – the furrow moves to the anterior while the posterior nucleus moves to the posterior (Fig.2F,G and movie 6). Opposite DNA and furrow movements thus allow correction of DNA segregation defects due to the relative mispositioning of the furrow and the mitotic spindle.

**Figure 2:**
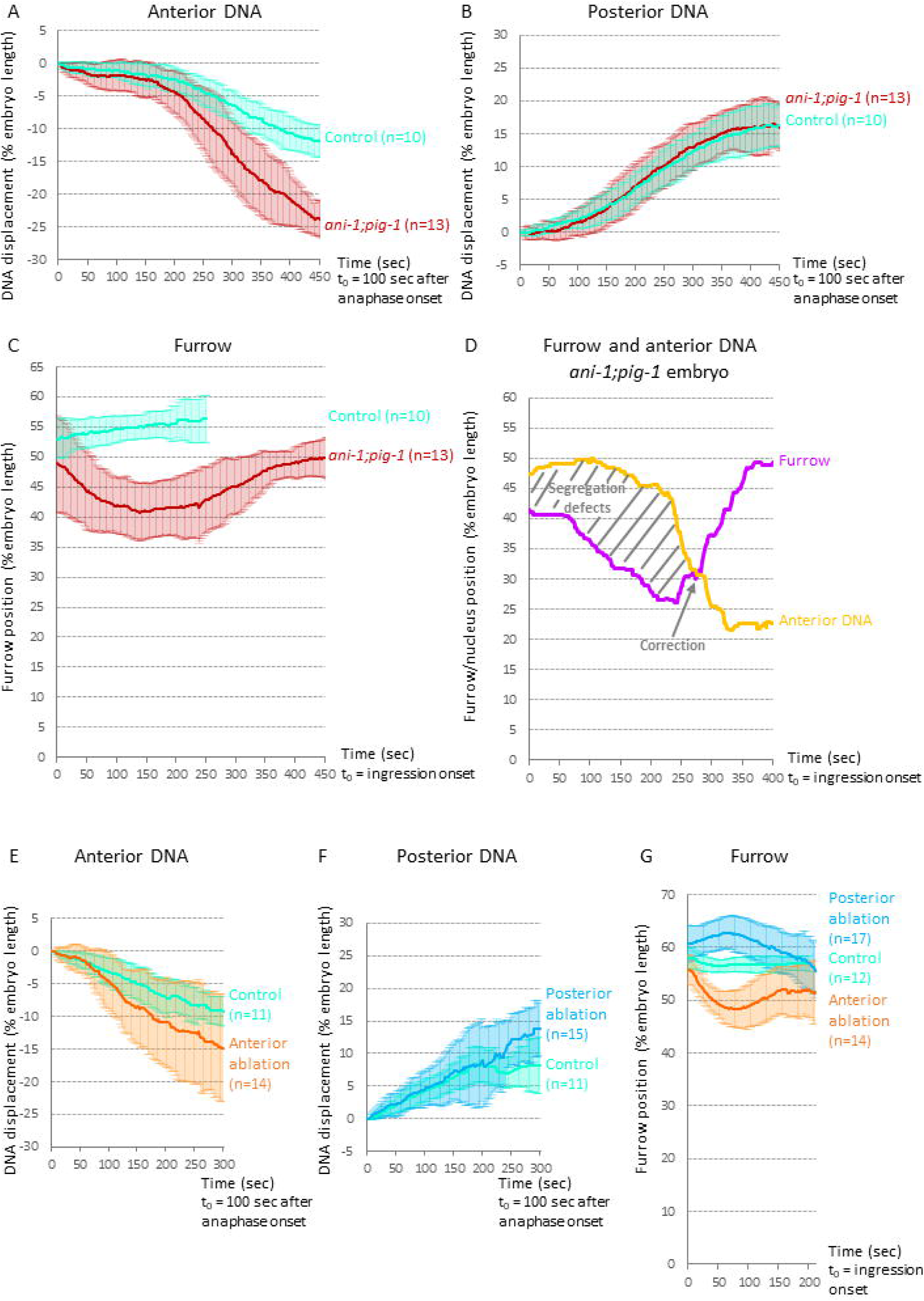
Correction of DNA segregation defects involves both nuclear and furrow displacement. **A-B.** Average displacement of the anterior (A) and posterior (B) nuclei in control and *ani-1(RNAi);pig-1(gm344)* embryos. Positive and negative values correspond to a displacement towards the posterior and anterior pole of the embryos, respectively. t_0_ = 100 sec after anaphase onset. **C.** Average furrow tip position along the antero-posterior axis in control and *ani-1(RNAi);pig-1(gm344)* embryos. 100 and 0 correspond to the posterior and anterior pole of the embryos, respectively. t_0_ = furrow ingression onset. **D.** Anterior nucleus and furrow tip position in a representative *ani-1(RNAi);pig-1(gm344)* embryo. 100 and 0 correspond to the posterior and anterior pole of the embryos, respectively. t_0_ = furrow ingression onset. In A-D, embryos recorded for those experiments express NMY-2::GFP and mCherry::HIS. **E-F.** Average displacement of the anterior (E) and posterior (F) nuclei in control embryos and in embryos in which the anterior (E) or posterior (F) centrosome was ablated during early anaphase. Positive and negative values correspond to a displacement towards the posterior and anterior pole of the embryos, respectively. t_0_ = 100 sec after anaphase onset. **G.** Average furrow tip position along the antero-posterior axis in control embryos and in embryos in which the anterior or posterior centrosome was ablated during early anaphase. 100 and 0 correspond to the posterior and anterior pole of the embryos, respectively. t_0_ = furrow ingression onset. In E-G, embryos express NMY-2::GFP, GFP::a-tubulin and GFP::HIS. In all panels, distances are expressed in percentage of embryo length and error bars correspond to standard deviation. Only embryos with corrected DNA segregation defects and complete furrow ingression were analysed.

### Nuclear microtubule interactions contribute to nuclear displacement

We next searched for the mechanisms involved in regulating nuclear displacement and first tested the role of microtubule/nuclear interactions. In the one-cell *C. elegans* embryo, microtubules were shown to control pronuclear position through their interaction with ZYG-12, a KASH protein located in the nuclear envelope. Thanks to a *zyg-12* thermosensitive allele *(zyg-12(or577)), we* inactivated ZYG-12 at anaphase onset to prevent the interaction between the nuclei and microtubules during the last steps of mitosis (see Key Resources table). In control and *ani-1;pig-1* embryos, interactions between nuclei and microtubules allow the close apposition of the centrosomes to the nuclei during late mitosis (Fig.3A). On the contrary, in *zyg-12* and *zyg-12;ani-1;pig-1* embryos, the centrosomes stay away from the nuclei (Fig.3A, see legend for quantifications), indicating that ZYG-12 inactivation efficiently prevents nuclei/microtubule interactions. Interestingly, ZYG-12 inactivation strongly inhibits anterior nuclear displacement in control embryos (Fig.3B). Nuclear/microtubule interactions are thus critical to move nuclei away from the cytokinetic furrow in control embryos. In *ani-1;pig-1* embryos, ZYG-12 inactivation nevertheless only moderately inhibits anterior nuclear displacement (Fig.3C) and does not affect the correction of DNA segregation defects (16/16 and 13/14 *ani-1;pig-1* and *zyg-12;ani-1;pig-1* embryos with corrected DNA segregation defects, respectively). Hence, nuclear/microtubule interactions are involved but not essential for anterior nuclear displacement in that context.

**Figure 3:**
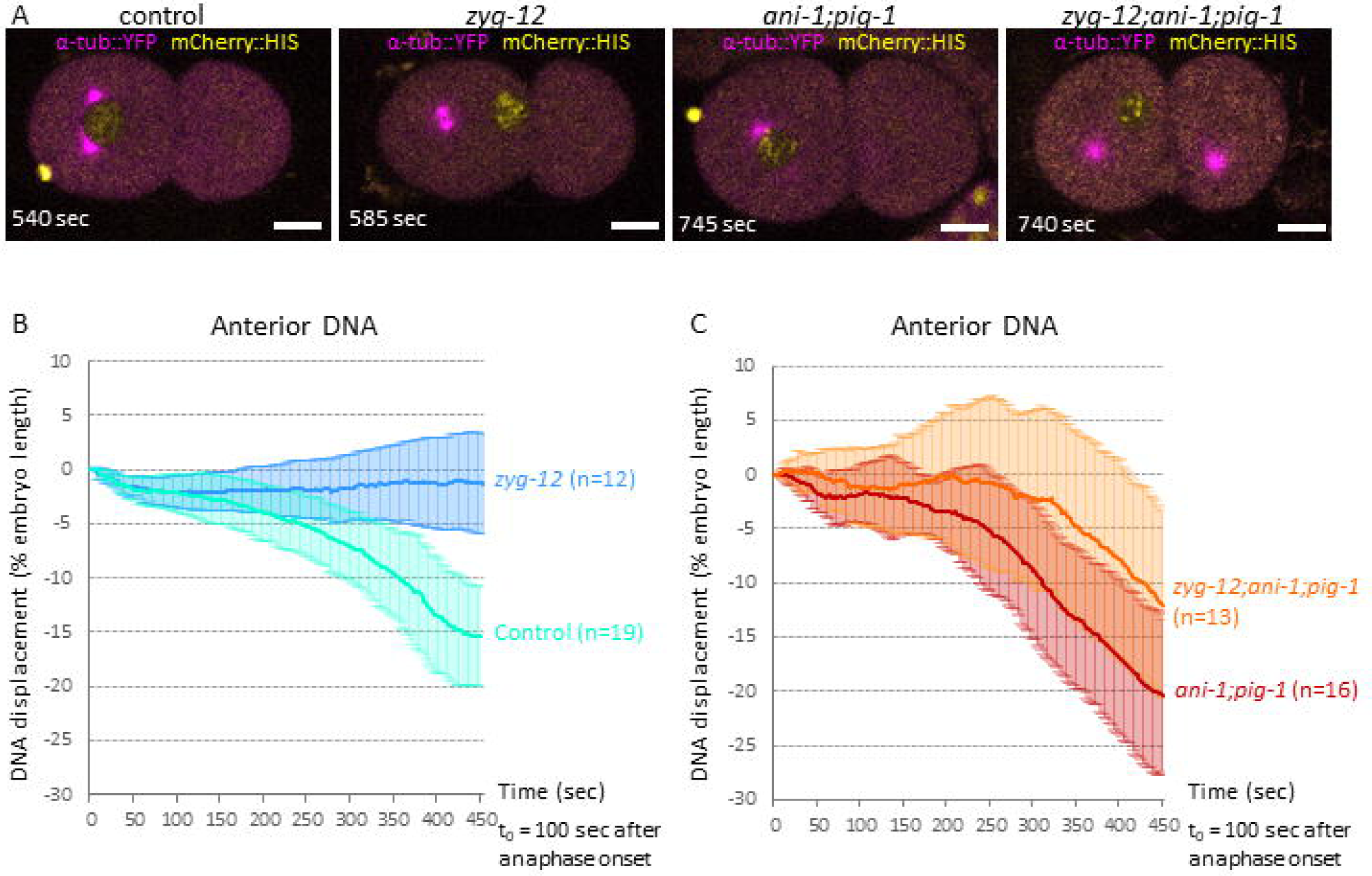
Nuclear microtubule interactions contribute to nuclear displacement. **A.** Control, *zyg-12(or577), ani-1(RNAi);pig-1(gm344)* and *zyg-12(or577);ani-1(RNAi);pig-1(gm344)* embryos expressing a-tubulin::YFP (magenta) and mCherry::HIS (yellow). In control (n=9) and *ani-1(RNAi);pig-1(gm344)* (n=12) embryos, the centrosomes are closely juxtaposed to the nuclei on average 435 seconds after anaphase onset. In *zyg-12(or577)* embryos (n=9), we never observed the centrosomes in close juxtaposition to the nuclei (embryos were observed until the duplication of centrosomes preceding the second division). In *zyg-12(or577), ani-1(RNAi);pig-1(gm344)* embryos (n=17), 14 embryos never showed juxtaposed centrosomes and nuclei while juxtaposition occurred very late (>500 sec after anaphase onset) in 3 embryos. In the embryos shown, posterior nuclei and most posterior centrosomes are out of focus. The second anterior centrosome is also out of focus in *ani-1;pig-1* and *zyg-12;ani-1;pig-1* embryos. t_0_ = anaphase onset. Embryos are oriented with the anterior to the left. Scale bar: 10 μm. **B-C.** Average displacement of the anterior nucleus in control, *zyg-12(or577), ani-1(RNAi);pig-1(gm344)* and *zyg-12(or577);ani-1(RNAi);pig-1(gm344)* embryos. All embryos express NMY-2::GFP and mCherry::HIS. Positive and negative values correspond to a displacement towards the posterior and anterior pole of the embryos, respectively; distances are expressed in percentage of embryo length. Error bars correspond to standard deviation. t_0_ = 100 sec after anaphase onset.

### Myosin activity regulates both nuclear and furrow displacement

We next looked for an additional mechanism that could participate in anterior nucleus displacement in *ani-1;pig-1* embryos and tested the possible involvement of myosin by using a thermosensitive allele of non-muscle myosin-2 *(nmy-2(1490)). nmy-2;ani-1;pig-1* embryos were initially grown at permissive temperature to ensure that the increase in myosin activity due to the lack of PIG-1 and ANl-1 is sufficient to induce some furrow mispositioning. They were then shifted at restrictive temperature 220 seconds after anaphase onset (Key Resources table). Myosin inactivation moderately reduces nuclear displacement in control embryos (Fig.4A). Thus, contrary to nuclear/microtubule interactions, myosin has only a minor role in the regulation of nuclear position in wildtype embryos. Myosin inactivation also moderately inhibits anterior nuclear displacement in *ani-1;pig-1* embryos (Fig.4B). Notably, simultaneous inactivation of ZYG-12 and myosin severely reduces the efficiency of DNA segregation defect correction. Indeed, while all DNA segregation defects are corrected in *nmy-2;ani-1;pig-1* embryos (12/12), only 55% (11/20) of *nmy-2;zyg-12;ani-1;pig-1* embryos with DNA segregation defects show a correction. Furthermore, the displacement of the anterior nucleus towards the anterior pole is strongly impaired, both in rescued and non-rescued *nmy-2;zyg-12;ani-1;pig-1* embryos (Fig.4C-C’). Altogether, our results demonstrate that microtubules are essential to generate the forces which regulate nuclear position at the end of division in control embryos while microtubules and myosin contribute separately to the positioning of the anterior nucleus during the correction of DNA segregation defects in *ani-1;pig-1* embryos.

**Figure 4:**
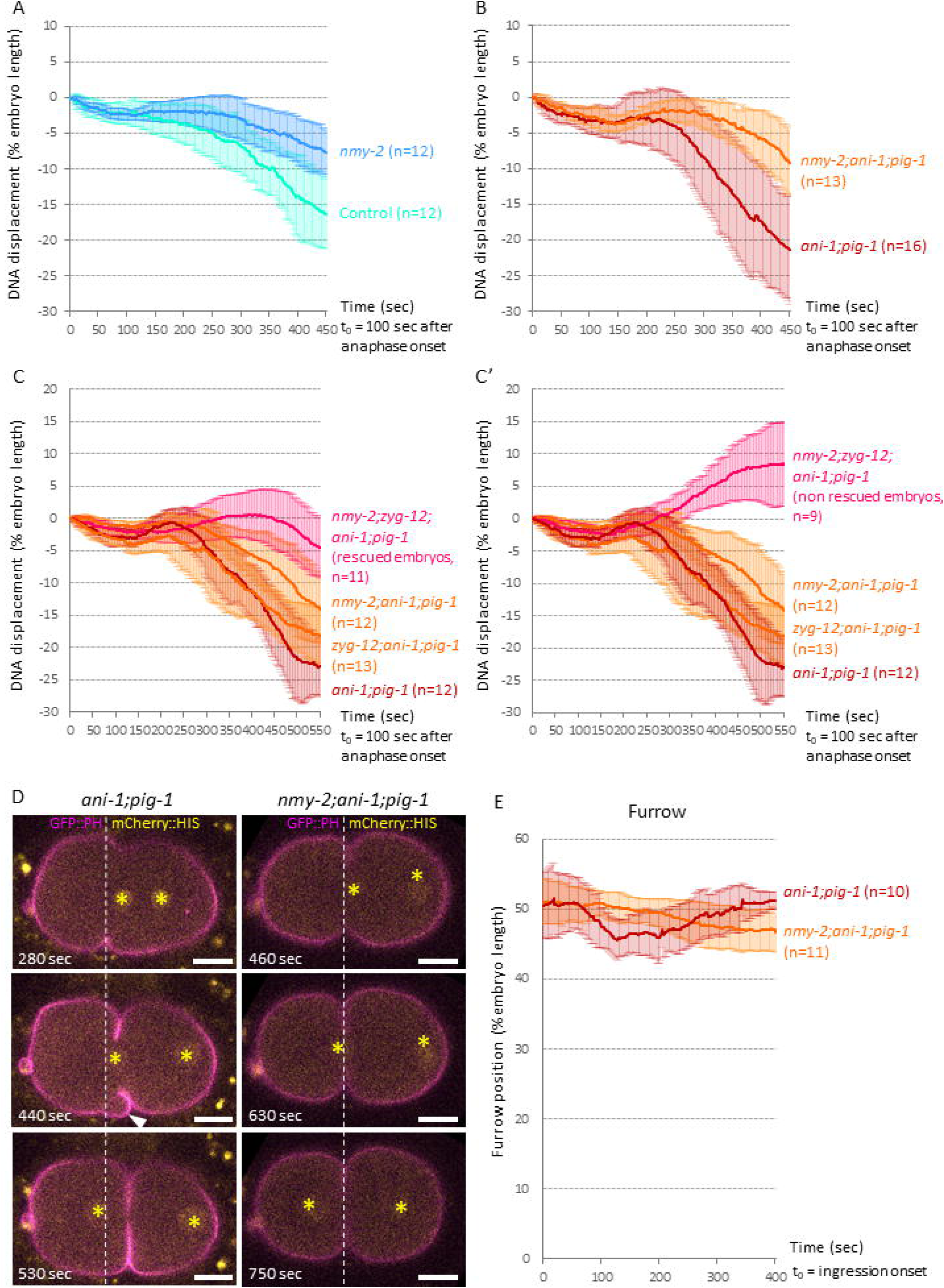
Myosin activity regulates both nuclear and furrow displacement. **A-B.** Average displacement of the anterior nucleus in control, *nmy-2(ne1490), ani-1(RNAi);pig-1(gm344)* and *nmy-2(ne1490);ani-1(RNAi);pig-1(gm344)* embryos. In B, only embryos with corrected DNA segregation defects were analysed. **C-C’.** Average displacement of the anterior nucleus in *ani-1(RNAi);pig-1(gm344), zyg-12(or577);ani-1(RNAi);pig-1(gm344), nmy-2(ne1490);ani-1(RNAi);pig-1(gm344)* and *nmy-2(ne1490);zyg-12(or577);ani-1(RNAi);pig-1(gm344)* embryos. Data presented in C and C’ were all obtained during the same experiment. For *ani-1(RNAi);pig-1(gm344), zyg-12(or577);ani-1(RNAi);pig-1(gm344)* and *nmy-2(ne1490);ani-1(RNAi);pig-1(gm344)*, only embryos with corrected DNA segregation defects were analysed (same set of embryos in C and C′). For *nmy-2(ne1490);zyg-12(or577);ani-1(RNAi);pig-1(gm344)*, embryos with corrected DNA segregation defects are presented in *C*, embryos without correction in C′. In A-C′, all embryos express mCherry::HIS; positive and negative values correspond to a displacement towards the posterior and anterior pole of the embryos, respectively; t_0_ = 100 sec after anaphase onset. **D.** *ani-1(RNAi);pig-1(gm344)* and *nmy-2(ne1490);ani-1(RNAi);pig-1(gm344)* embryos expressing GFP::PH (magenta) and mCherry::HIS (yellow, DNA also marked with yellow asterix). White dashed line indicate the most anterior position reached by the furrow. The furrow is then displaced towards the posterior in *ani-1(RNAi);pig-1(gm344)* embryos but not in *nmy-2(ne1490);ani-1(RNAi);pig-1(gm344)* embryos. In *ani-1(RNAi);pig-1(gm344)* embryos, furrow displacement is associated with the formation of cortical blebs on the anterior side of the furrow (white arrowhead, 10/10 embryos). Such blebs are never observed in *nmy-2(ne1490);ani-1(RNAi);pig-1(gm344)* embryos (n=11). t_0_ = anaphase onset. Embryos are oriented with the anterior to the left. **E.** Average furrow tip position along the antero-posterior axis in *ani-1(RNAi);pig-1(gm344)* and *nmy-2(ne1490);ani-1(RNAi);pig-1(gm344)* embryos expressing GFP::PH and mCherry::HIS. Only embryos with corrected DNA segregation defects were analysed. 100 and 0 correspond to the posterior and anterior pole of the embryos, respectively; t_0_ = furrow ingression onset. In all panels, distances are expressed in percentage of embryo length and error bars correspond to standard deviation.

Importantly, we also found that myosin inactivation prevents the furrow from moving back towards the posterior in *nmy-2;ani-1;pig-1* embryos (Fig.4D-E, movie 7). Myosin therefore plays a crucial role in the correction process by concomitantly regulating nuclear and furrow displacements.

### Variations of myosin levels entail changes in cortical tension

Considering the importance of myosin in the correction process, we then carefully examined myosin cortical levels. As previously described, the anterior cortical level of myosin briefly increases in control anaphase embryos while an excessive and long-lasting accumulation of myosin is observed in *ani-1;pig-1* embryos (Fig.5A, left) (Pacquelet et al., 2015). This accumulation then slowly drops off during the correction process (Fig.5A, left). Those changes in myosin levels are specific to the anterior cortex, as posterior cortical levels of myosin increase only weakly and briefly in anaphase control and *ani-1;pig-1* embryos (Fig.5A, right). Those observations prompted us to assess possible changes in cortical tension. To this end, we performed laser-induced cortical ablation (Mayer et al., 2010; Saha et al., 2016) and measured the velocity of laser-induced cortical cut opening both in control and *ani-1;pig-1* embryos (Fig.5B; movie 8; see STAR methods for details). In control embryos, cortical tension during anaphase is slightly higher at the anterior than at the posterior cortex (Fig.5C). While posterior cortical tension does not change significantly in *ani-1;pig-1* embryos compared to control embryos, it strongly increases at the anterior cortex during anaphase before declining at the end of the rescue process (Fig.5C). All these variations in cortical tension are consistent with the changes in myosin levels that we measured (Fig.5A).

**Figure 5:**
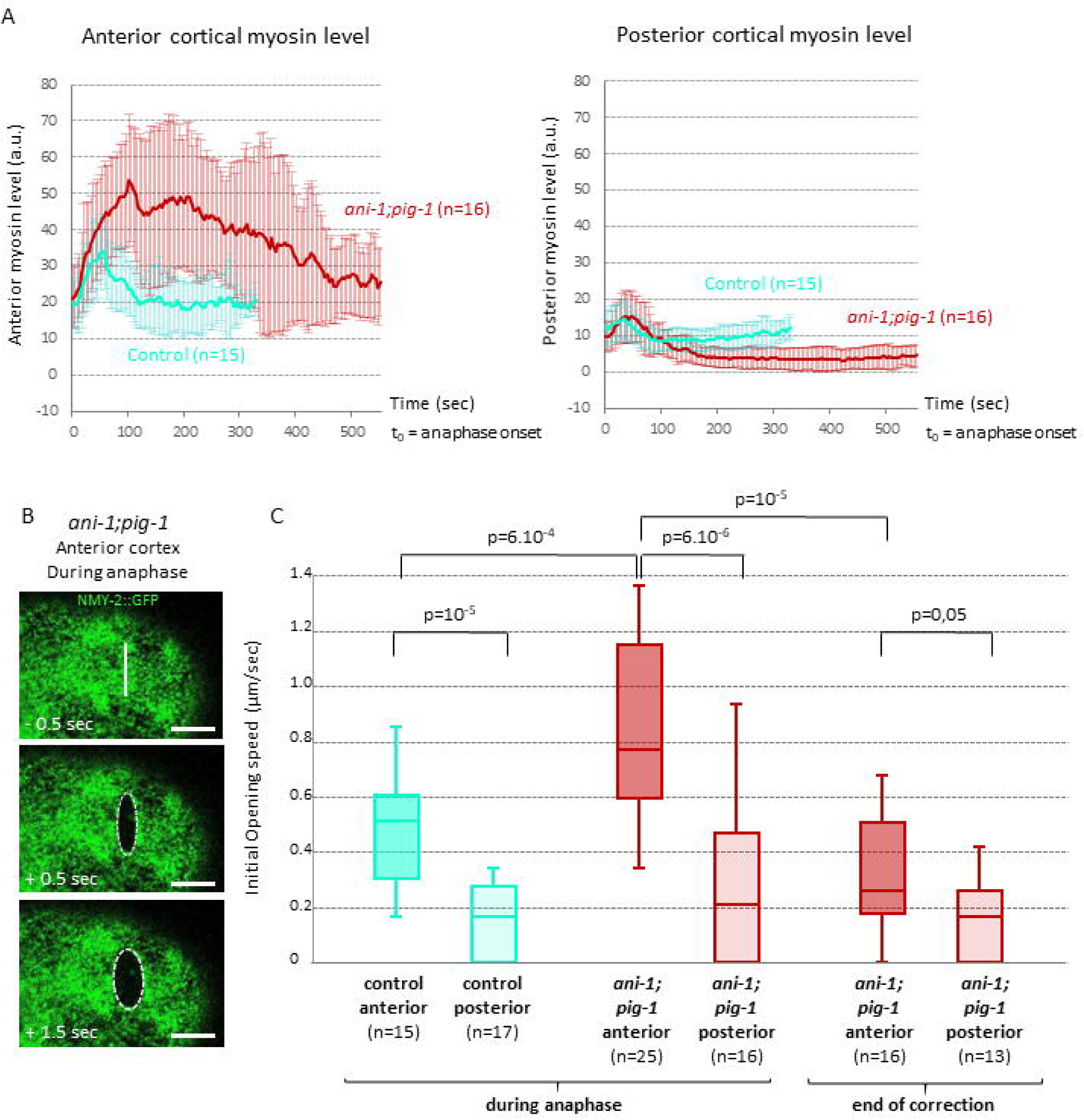
Variations in myosin concentration entail different levels of cortical tension. **A.** Anterior (left) and posterior (right) cortical myosin levels in control and *ani-1(RNAi);pig-1(gm344)* embryos expressing NMY-2::GFP and mCherry::HIS. t_0_ = anaphase onset. Error bars represent standard deviation. **B.** Laser-induced ablation performed during anaphase at the anterior cortex of a *ani-1(RNAi);pig-1(gm344)* embryo. Images of the anterior cortex were taken 0.5 sec before, 0.5 sec and 1.5 sec after ablation. White bar indicates the area targeted for laser ablation. Measurement of the ellipse small axis allowed us to determine the cut opening kinetics (see STAR methods for details). Scale bar: 5 µm. **C.** Quantification of initial opening velocities at the anterior and posterior cortex of control and *ani-1(RNAi);pig-1(gm344)* embryos, p-values: Wilcoxon test. In all panels, embryos express NMY-2::GFP (green) and mCherry::HIS (not shown).

### Changes in cortical myosin levels contribute to the repositioning of the nascent furrow

We next asked whether the changes in anterior cortical myosin levels and cortical tension that we observed could affect furrow position. It was indeed previously shown that cortical tension imbalance and asymmetric myosin-driven contraction of the cortex can alter daughter cell size in dividing Hela cells (Sedzinski et al., 2011). This suggested that the variations in myosin level and cortical tension that we observe in *ani-1;pig-1* embryos could explain furrow position movements. Consistent with this hypothesis, we found that furrow movement is correlated with cortical myosin levels: large differences of myosin levels between the anterior and posterior cortex (Δmyosin > 50 a.u.) are associated with the anterior displacement of the nascent furrow while weaker differences (Δmyosin < 30 a.u.) coincide with the furrow moving back towards the posterior pole of the embryo (Fig.6A).

**Figure 6:**
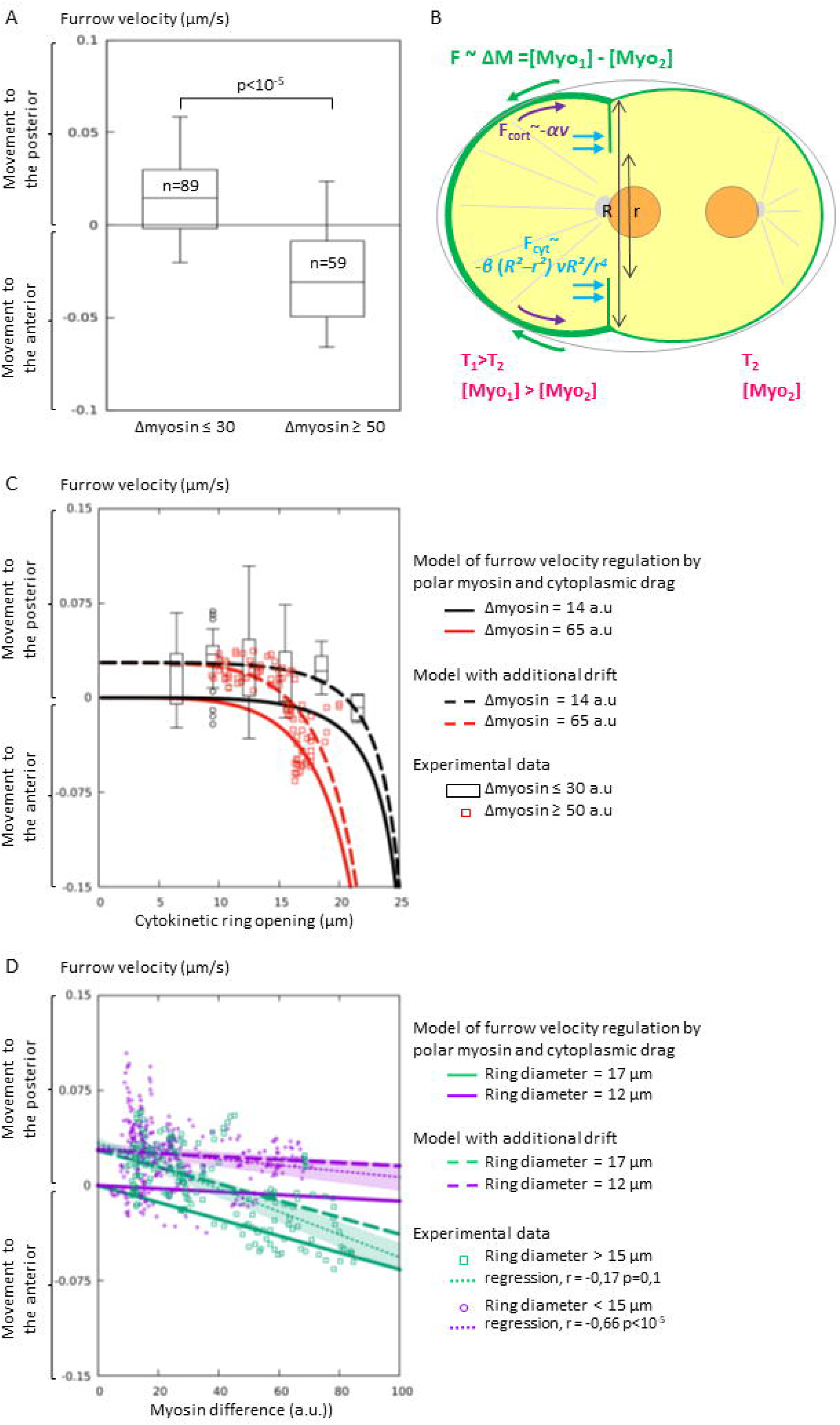
Cortical myosin can reposition the nascent furrow. A. Furrow velocity of nascent cytokinetic furrows (ring diameter > 15 µm) depends on ΔM (= difference between anterior and posterior myosin levels), p-value: Student t-test. **B.** Model of the forces acting on furrow displacement. The stronger contractility of the anterior cortex exerts a pulling force on the nascent cytokinetic ring (F ∼ ΔM) while resistance forces from the cortex (F_cort_) and the cytoplasm (_Fcyt_) oppose to furrow displacement. *R* corresponds to the cell radius and *r* to the cytokinetic ring radius. **C.** Furrow velocity as a function of ring opening and myosin concentration. Black boxplots and red squares measurements correspond to low and high ΔM, respectively (n=89 for low ΔM and n=59 for high ΔM, time points collected from 15 different embryos, see STAR methods). As detailed in the STAR methods section, modelling the forces exerted by myosin and the resistance forces predicts that furrow velocity scales as *v* ∼ ΔM *r*^4^ /(α*r*^*4*^ ¿ *ß R*^*2*^*(R*^*2*^*-r*^*2*^)). The predicted velocity for low and high ΔM is represented by the solid black and red lines, respectively. A constant drift velocity needs to be added to the model (black and red dashed lines) to fit the experimental data. **D.** Furrow velocity as a function of myosin concentration and ring opening. Measured furrow velocity is correlated with ΔM in the case of nascent furrow (ring diameter > 15 pm, green squares and green dotted regression line, | r | =0.66, p<10^-5^) but not during ring closure (ring diameter < 15 pm, purple dots and purple dotted regression line | r | =0.17, p=0.1). Furrow velocity predicted by our model is represented without (solid lines) or with (dashed lines) addition of a constant drift, for a nascent furrow (ring diameter = 17 µm, green) or during ring closure (ring diameter = 12 µm, purple). In all panels, furrow velocity was measured between 300 sec and 450 sec after anaphase onset (see STAR methods for details). Positive and negative values of furrow velocity correspond to a displacement towards the posterior and anterior pole of the embryos, respectively.

If this correlation results from the mechanical action of cortical myosin on the furrow, we reasoned that furrow speed must be affected by factors that influence the mechanical resistance to furrow movements. Forces resisting furrow displacement include the resistance of the cortex itself, which can be assumed proportional to furrow velocity, *F*_*cort*_ = −αv (Fig.6B). It also involves the drag force exerted by the cytoplasm, which, due to the confinement of the embryo in its eggshell, needs to flow through the cytokinetic ring in order to allow displacement of the furrow in the opposite direction (Fig.6B). This latter resisting force is expected to increase when the cytokinetic ring closes. Specifically, assuming a Poiseuille flow through the cytokinetic ring, we expect a dynamic pressure drop between the two daughter cells scaling as *vR*^*2*^*/r*^*4*^, where *R* is the cell radius and *r* the cytokinetic ring radius. Then, the drag force exerted by the cytoplasm will be *F*_cyt_ = *ß (R*^*2*^*−r*^*2*^*) vR*^*2*^*/r*^*4*^. If the difference of cortical myosin concentration between anterior and posterior, ΔM, drives furrow motion, we expect the velocity of the furrow to scale as *v ∼ ΔM r*^*4*^ */(αr*^*4*^ *+ ß R*^*2*^*(R*^*2*^*-r*^*2*^)) (see STAR methods for details). This model predicts that for a given myosin difference furrow speed depends on cytokinetic ring closure (Fig.6C, red continuous line (Δmyosin > 50 a.u)). Moreover, the effect of myosin concentration imbalance will become negligible as the furrow closes (Fig.6C, compare red (Δmyosin > 50 a.u.) and black (Δmyosin < 30 a.u.) continuous lines and Fig.6D, compare green (ring diameter > 15 μm) and purple (ring diameter < 15 μm) continuous lines). In agreement with this model, we found that for a given myosin difference furrow velocity strongly depends on cytokinetic ring opening (Fig.6C, red squares (Δmyosin > 50 a.u.)). We also observed that the speed of the nascent furrow is correlated with myosin levels (Fig.6C, ring diameter > 15 μm, green squares and green dotted regression line) while this is not the case when the furrow ingresses deeper into the cytoplasm (Fig.6D, ring diameter < 15 μm, purple squares and purple dotted regression line). Altogether, our data strongly suggest that changes in myosin cortical levels contribute to the displacement of the nascent cytokinetic furrow. However, as the cytokinetic ring closes, furrow movement is limited by the increasing cytoplasmic drag, suggesting the existence of a supplementary mechanism that contributes to further furrow displacement. Moreover, consistent with the hypothesis that changes in myosin levels is not the only mechanism driving furrow displacement, we observed a shift between the velocity of the furrow predicted by our model and our experimental data (Fig.6C-D). In particular, while the model predicts that myosin influences the speed of the furrow movement towards the anterior cortex, it does not account for the reverse movement of the furrow towards the posterior (Fig.6C-D, continuous lines) and a constant drift velocity must be added to our model to fit experimental data (Fig.6C-D, dashed lines, see STAR methods for details).

### Myosin-dependent cortical blebbing and cytoplasmic flow are associated with furrow displacement

We thus searched for additional mechanisms that could explain furrow repositioning. Displacement of the furrow towards the posterior could conceivably follow the regulation of myosin activity by the central spindle. To assess the role of the central spindle, we used a thermosensitive allele to inactivate the central spindle protein SPD-1 during anaphase (see Key Resources table). This inactivation leads to the acceleration of anaphase chromosome segregation, indicating that it effectively disrupts the central spindle (Fig.S1A). In *ani-1;pig-1* embryos, it nevertheless does inhibit neither correction efficiency nor furrow displacement (Fig.S1B and legend, movie 9), showing that the central spindle does not play an essential role in repositioning the cytokinetic furrow. This is consistent with our previous observations showing that the correction of DNA segregation defects occurs late during the cell cycle, when the central spindle has already started to disassemble (Fig.1C, movie 4). Alternatively, the abnormal localization of the anterior nucleus relative to the cytokinetic furrow could contribute to furrow repositioning. Indeed, previous work in *Drosophila* has shown that nuclei can regulate cortical cell shape changes by sequestering myosin regulators such as Ect2 (Montembault et al., 2017). However, preventing nuclear reformation by depleting the nucleoporin NPP-8 (Fig.S1C) does not prevent correction of DNA segregation defects or furrow displacement (Fig.S1D and legend, movie 10), indicating that the nuclei do not control furrow repositioning through the sequestration of myosin regulators.

While none of the signalling pathways that we tested appears to control furrow position, our observations suggest a possible involvement of cortical blebbing and cytoplasmic flow, especially during cytokinetic ring closure. Indeed, *ani-1;pig-1* embryos display strong cortical deformations, which are not seen in control embryos. Those deformations are characterized by a sudden local decrease in cortical myosin levels (Fig.7A, movie 11) and are very similar to the blebs that have typically been described in cultured cells (Charras et al., 2008). Cortical blebs first arise at the posterior and anterior embryo poles (Fig.7A, movie 11). They form early during anaphase, prior to or at the beginning of cytokinesis furrow ingression, when the furrow still moves towards the anterior pole of the embryo (Fig.7A-B, movie 11). About 250 seconds after anaphase onset, those posterior and anterior blebs cease and are replaced by cortical deformations located on the anterior edge of the cytokinetic furrow (Fig.7A-B and movie 11). The blebs of this latter type form during furrow ingression, appearing shortly prior to and then accompanying furrow posterior displacement (Fig.7A-B, movie 11). Importantly, they do not form when myosin is inactivated (Fig.4D, movie 7, see figure legend for quantifications). These results suggest that bleb formation is likely due to the increased cortical tension that was measured in *ani-1;pig-1* embryos (Fig.5C), similar to what was previously shown in cultured cells (Charras et al., 2008).

**Figure 7:**
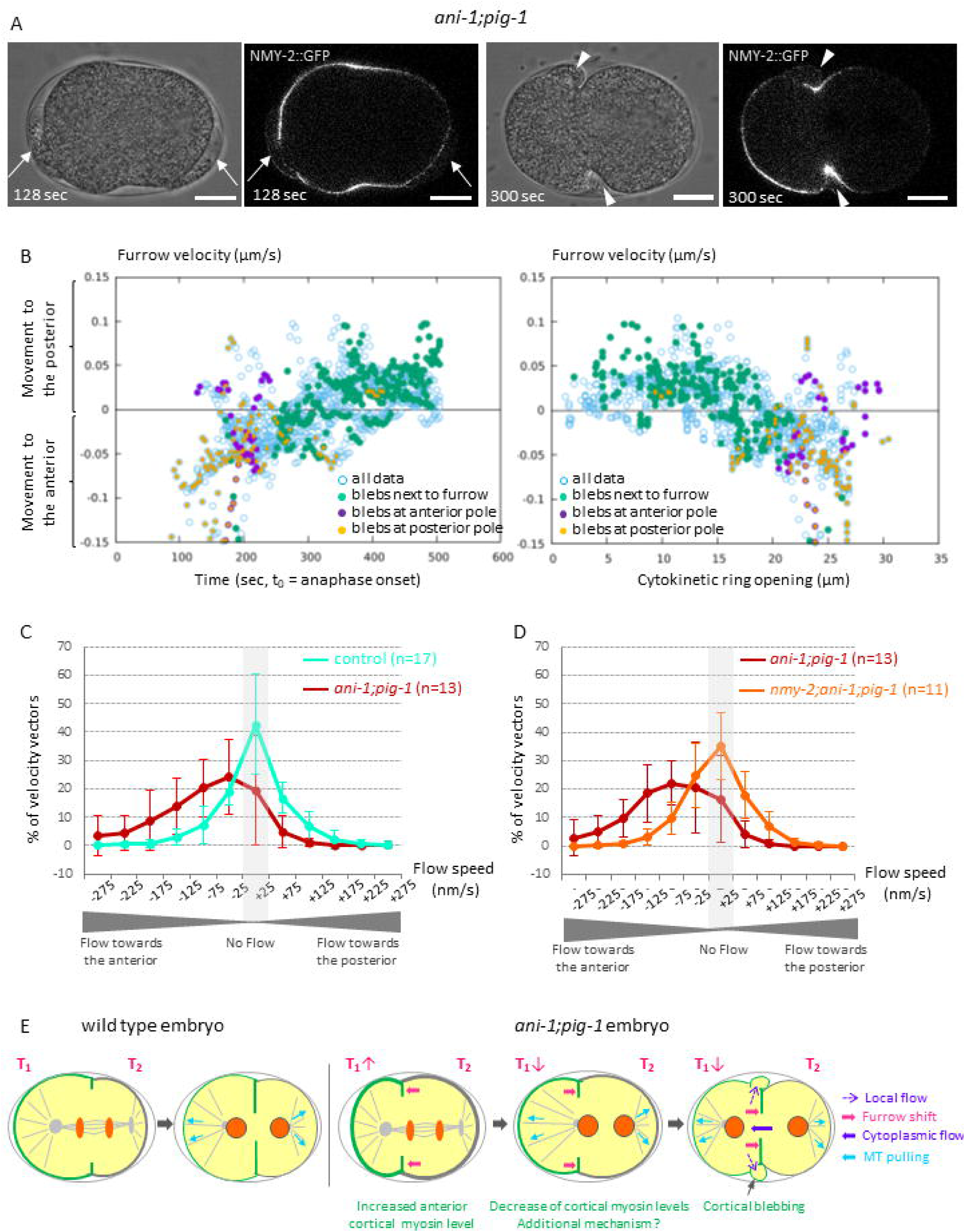
Furrow repositioning is accompanied by cortical blebbing and cytoplasmic flow. **A.** Formation of cortical blebs on the anterior and posterior poles (t=128 sec, arrows) and on the anterior side of the cytokinetic furrow (t=300 sec, arrowheads) in a *ani-1(RNAi);pig-1(gm344)* embryo expressing NMY-2::GFP (gray) and mCherry::HiS (not shown). The cortical blebs identified on DIC images (left) are characterized by a transient decrease in cortical myosin level (images on the right), to = anaphase onset. Embryo is oriented with the anterior to the left. Scale bar: 10 µm. **B.** Presence of blebs at the posterior (orange dots) and anterior (purple dots) poles of the embryos precede the formation of blebs close to the furrow (green dots). Blue circles represent all time points. Time points were collected from 15 different embryos (see STAR methods). Positive and negative values of furrow velocity correspond to a displacement towards the posterior and anterior pole of the embryos, respectively. t_0_ = anaphase onset. **C-D.** Distribution of velocity vector intensities measured by PIV in control, *ani-1(RNAi);pig-1(gm344)* and *nmy-2(ne1490);ani1(RNAi);pig-1(gm344)* embryos expressing vit-2::GFP and mCherry::HIS. Flow speed was measured along the antero-posterior axis, positive and negative values correspond to a flow towards the posterior and anterior pole of the embryo, respectively. Only embryos with corrected DNA segregation defects were analysed. Error bars correspond to standard deviation. **E.** Model of the mechanisms driving furrow and nuclear positions in wildtype and *ani-1(RNAi);pig-1 (gm344)* embryos. In wildtype embryos, anterior cortical myosin level is downregulated during division. The differential contractility between the anterior and posterior cortices remains low and is therefore not sufficient to displace significantly the furrow from the location imposed by mitotic spindle signalling. In *ani-1(RNAi);pig-1(gm344)* embryos, the strong increase of anterior myosin levels in anaphase exerts a pulling force on the nascent cytokinetic ring, moving it towards the anterior cortex. The subsequent decrease of anterior myosin arrests this movement. Once the furrow closes, cortical blebbing generates a pressure decrease in the anterior daughter cell. The ensuing cytoplasmic flow then drives the opposite displacement of the cytokinetic furrow and the anterior nucleus. Microtubules also contribute to the correction process by pulling on the anterior nucleus.

Blebbing can transiently reduce intracellular pressure (Tinevez et al., 2009). This suggests that when the cytokinetic ring is sufficiently narrow, bleb formation on the anterior side of the furrow could induce a pressure difference between the two daughter cells. Such a pressure difference would in turn drive a cytoplasmic flow. Consistent with this hypothesis, we observed that posterior furrow displacement also coincides with the appearance of a strong flow of cytoplasmic particles, which is oriented towards the anterior pole of *ani-1;pig-1* embryos (movie 12). We used Particle Imaging Velocimetry (PIV) to quantify this flow and found that 75% of the measured velocity vectors are oriented towards the anterior in *ani-1;pig-1* embryos (speed<-25nm/s, n=13 embryos, Fig.7C). Such a flow is not observed in control embryos (only 30% of the velocity vectors with speed<-25nm/s, n=17, movie 11, Fig.7C) or when myosin is inactivated (39% of the velocity vectors with speed<-25nm/s, n=11, Fig.7D). Similarly, the correction of DNA segregation defects following anterior centrosome ablation involves both the displacement of the anterior nucleus and of cytoplasmic particles towards the anterior pole (movie 13). Reversely, the correction of defects due to posterior centrosome ablation is associated with the displacement of both the posterior nucleus and cytoplasmic particles towards the posterior (movie 13).

Altogether, our results strongly suggest that formation of cortical blebs on the anterior side of the furrow locally releases pressure and induces a cytoplasmic flow towards low pressure areas. This effect is expected to be particularly important when the cytokinetic ring closes (see discussion for further details). Since the embryo is confined in its eggshell, the flow of cytoplasmic material towards the anterior must then be compensated through the repositioning of the furrow towards the posterior pole of the embryo. It is thus very likely that cortical blebbing and the subsequent cytoplasmic flow contribute to furrow repositioning during ring closure (Fig.7E).

## Discussion

Altogether, our work shows that DNA segregation defects due to cytokinetic furrow mispositioning in one-cell *C. elegans* embryos can be corrected during late mitosis. This so far uncharacterized correction mechanism relies on both cytokinetic furrow and nucleus displacement. It is also accompanied by a strong cytoplasmic flow and by blebbing of the cortex adjacent to the furrow. Importantly, we found that correction of DNA segregation defects due to furrow mispositioning is not specific to *ani-1;pig-1* embryos. It is for instance also observed when relative spindle / furrow mispositioning is induced upon centrosome ablation. Similar to what happens in *ani-1;pig-1* embryos, correction of DNA segregation defects in embryos with ablated centrosomes also involve opposite DNA and furrow movements as well as cytoplasmic flow. It is therefore very likely that comparable mechanisms are at play in both cases.

We have shown that myosin plays a crucial role in the regulation of the different facets of the correction process. First, we find that the difference between anterior and posterior myosin concentrations control the displacement of the nascent cytokinetic furrow. The different contractility of the anterior and posterior cortices can indeed exert a pulling force on the nascent cytokinetic ring, oriented towards the more contractile end. In control embryos, this differential contractility remains low and is therefore not sufficient to displace significantly the furrow from the location imposed by mitotic spindle signalling (Fig.7E). On the contrary, the strong increase of anterior myosin levels in anaphase *ani-1;pig-1* embryos can explain the initial shift of the nascent furrow towards the anterior cortex. The subsequent decrease of anterior myosin then slows down this movement. It can however not explain the repositioning of the furrow towards the posterior. Moreover, we show that differential cortical contractions do not participate in driving furrow displacement during the whole process of DNA segregation defect correction. Indeed, as the ring closes, the posteriorly moving furrow behaves like a mechanical piston, which needs to displace cytoplasm anteriorly to allow its own motion. Our theoretical model indicates that this creates a drag force, which is too large to be overcome by cortical contractility.

Our observations strongly suggest that the blebs forming on the anterior edge of the furrow as well as the strong cytoplasmic flow directed towards the anterior pole of the embryo contribute to furrow displacement during cytokinetic ring closure. Highly contractile cortices favour the appearance of cortical blebbing in *ani-1;pig-1* embryos. In those embryos, blebs appear first predominantly at the anterior or posterior poles and then close to the furrow once this latter has formed. Notably, these locations correspond to the regions where the cortex is away from the eggshell and with the strongest cortical curvature (Fig.7A). Following the theory of bleb nucleation proposed in (Charras et al, 2008), we modelled the variation of free energy entailed by the nucleation of a bleb in the different regions of an embryo. In agreement with our observations, this model predicts a higher probability of forming a bleb where the cortex is strongly curved (for details see model in STAR methods). Interestingly, blebbing has been shown to transiently reduce intracellular pressure (Tinevez et al., 2009). We therefore suggest that the cytoplasmic flow that we observe is a consequence of the “depressurizing effect” of cortical blebs: as they form on the anterior side of the furrow, the anterior daughter cell pressure decreases; because of the narrow cytokinetic ring, pressure equilibration with the posterior daughter cell is comparatively slow and their transient pressure difference drives a flow from the posterior cell to the anterior cell. Due to embryo confinement, the resulting increase in the volume of the anterior cell can then be accommodated only through the repositioning of the furrow to a more posterior location (Fig.7E). Altogether, our data are consistent with a model in which changes in anterior cortical myosin level contribute to movements of the nascent furrow while cortical blebs and subsequent cytoplasmic flow displace the furrow towards the posterior pole of the embryo when the cytokinetic furrow closes. It is however worth noting that changes in myosin levels are also not sufficient to explain furrow displacement towards the posterior in the early steps of furrow ingression. Cortical blebs might already weakly contribute to furrow displacement at this stage. Alternatively, a yet unknown biochemical signal may be involved. Our data show that the central spindle is not directly controlling furrow repositioning. Whether signalling at the cell equator or emanating from the embryo poles is involved remains to be determined.

Moreover, we found that myosin also regulates nuclear displacement. Notably, the anteriorly-directed movement of the nucleus and the flow of cytoplasmic particles occur simultaneously in *ani-1;pig-1* embryos. It is thus very likely that the cytoplasmic flow exerts a drag force on the anterior DNA to drive it through the cytokinetic ring. Hence, through its effect on cortical tension and deformations, myosin induces a strong cytoplasmic flow, which in turn regulates the opposite movements of the cytokinetic furrow and of the anterior nucleus. Finally, a supplementary and redundant mechanism based on nuclear/microtubule interactions also drives nuclear displacement – probably through the pulling of the nucleus by microtubules. The existence of these two mechanisms controlling nuclear displacement together with the coordination of furrow and nuclear displacements by the cytoplasmic flow are likely to increase the efficiency of DNA segregation defect correction (Fig.7E).

Our findings are reminiscent of what has previously been described in unconfined cultured cells (Sedzinski et al., 2011). Indeed, we find that excessive polar myosin contractility favours the risk of aneuploidy by mispositioning the cytokinetic furrow while cortical blebbing, which occurs in response to excessive cortical tension, contributes to the repositioning of the furrow, thereby helping to restore euploidy. In isolated cultured cells, variations in cortical tension induce changes in the Laplace force, which play a central role in cell shape regulation (Sedzinski et al., 2011). However, contrary to cultured cells, the one-cell *C. elegans* embryo is confined in a rigid eggshell. This restrains changes in cell curvature and creates a reaction force, which buffers changes in the Laplace force. Confinement thus creates geometrical constraints, which modulate the effects of cortical tension and the mechanisms involved in furrow repositioning. Moreover, our work highlights a new role for cortical blebs. While previous work suggested that blebs help providing more cortical surface and relaxing excessive pressure to stabilize cell shape during cell division (Sedzinski et al., 2011), we find that they can also regulate the volume of each daughter cell when the dividing cell is in a confined environment.

In conclusion, our work shows that dividing cells in a confined environment, as are most cells inside tissues, can benefit from a combination of rescue mechanisms, which prevents the polar actomyosin-driven destabilization of cytokinesis and can restore euploidy.

## Acknowledgments

We would like to thank P Askjaer, M. Delattre, J. Pécreaux and the *Caenorhabitis* Genetics Center (funded by the National Institute of Health Office of Research Infrastructure Programs, P40 OD010440) for providing worm strains. We also thank Wallis Nahaboo and M. Delattre for sharing unpublished observations. M. Ponnavoy helped with the earlier steps of this project. N. Loyer provided a macro for furrow and nuclear tracking and S. Prigent (Biogenouest, Université de Rennes 1) wrote a macro for PIV analysis. S. Dutertre and X. Pinson from the Microscopy Rennes Imaging Center (Biosit UMS 3480) assisted us with microscopy. S. Huet and Mathieu Pinot advised us for laser ablation experiments and Y Le Cunff for statistical analysis. We would also like to thank K. John (LIPhy) and members of G. Michaux and J. Pécreaux labs for discussions as well as G. Charras for critical reading of the manuscript.

This work was supported by institutional funding from the Centre National de la Recherche Scientifique (CNRS) and the Université Rennes 1. The group from G. Michaux is also supported by the Ligue Régionale contre le Cancer – Grand Ouest. J. Etienne is supported by the French National Research Agency in the framework of the “Investissements d’avenir” program (ANR-15-IDEX-02, IRS “AnisoTiss”), acknowledges CNRS Momentum grant “Modeling of living systems”, Tec21 (ANR-11-LABX-0030), and is member of GDR 3570 MécaBio and GDR 3070 CellTiss of CNRS.

## STAR Methods

### Contact for reagents and resource sharing

Further information and requests for resources and reagents should be directed to and will be fulfilled by the Lead Contact, Anne Pacquelet.

### Experimental model and subject details

#### Worm strains and culture

C. *elegans* strains were grown on agar plates containing NGM growth media and seeded with *E. coli* (OP50 strain). Worms were maintained at 20°C, except strains carrying thermosensitive mutations which were kept at 15°C. A list of the strains used in this study and conditions used for each experiment are detailed in the Key Resources table.

### Method details

#### RNAi conditions

RNAi was performed by feeding worms with RNAi clones from the Ahringer-Source Bioscience library (Kamath et al., 2003) on 1 mM or 3 mM IPTG plates. L4440 was used as a control in all RNAi experiments. The clone used for *rga-3/4* RNAi is primarily directed against *rga-3* but also targets *rga-4* (Schmutz et al., 2007). In the case of double RNAi experiments, RNAi cultures were initially grown separately; *ani-1* RNAi was then mixed either with control (L4440) or *npp-8* cultures (1:1) before seeding plates. Conditions used for each experiment are detailed in the Key Resources table.

#### Time-lapse recordings

Embryos were mounted on 2% agarose pads in a drop of M9 medium. In most experiments, temperature was controlled using a temperature-control platform, which enables temperature shifts within 10 seconds (CherryTemp, Cherry Biotech). Conditions used for each experiment are detailed in the Key Resources table.

#### Centrosome ablations

Centrosome laser ablations were performed on a Airy Scan microscope (Zeiss) with a 355 nm pulsed-UV laser (Rapp Optoelectronic, 40% laser + ND1 filter, 10 iterations / ablation) and a 40× water-objective. A circular area encompassing the anterior or posterior centrosome was defined for each ablation. Ablation was performed during early anaphase. Images were recorded every 2 sec.

#### Cortical ablations

Cortical laser ablations were performed on a Airy Scan microscope (Zeiss) with a 800 nm biphoton laser (Mai Tai, 50% power, 10 iterations / ablation) and a 63× oil-objective. A 0.17 × 6 µm rectangle perpendicular to the antero-posterior axis was used to define the cut area. The cut area was positioned on the anterior or posterior cortex, approximately at equal distance from the pole and the cytokinetic furrow. Ablations were performed during anaphase (1 min 10 sec to 1 min 50 sec after anaphase onset) or at the end of the correction process (5 to 8 min after anaphase onset). Images were recorded every 0.5 sec.

### Quantifications and statistical analysis

#### Image analysis and quantifications

Images were assembled for illustration using ImageJ. Position of the nuclei and of the furrow tip during ingression was tracked manually in ImageJ. Furrow tip position was measured starting at furrow ingression onset. Nuclei position tracking was generally started 100 sec after anaphase onset, when DNA separation due to anaphase spindle elongation is mostly finished. In Fig.S1A, we started to measure anterior nucleus displacement at anaphase onset to be able to evaluate the effect of central spindle disassembly on DNA movement during anaphase. Variations in experimental settings, in particular temperature shifts (see Key Resources table), explain the slight variations of nuclei or furrow dynamics observed between experiments for the same genotype.

Quantifications of myosin intensities (Fig.5A) were performed using ImageJ. A region corresponding to the most anterior or posterior 10–15% of the embryo cortex was defined and NMY-2::GFP cortical intensity was measured in a central confocal plane from anaphase onset to cytokinesis ring closure.

Furrow speed (Fig.6 and Fig.7B) was quantified based on furrow position by a LOESS regression over 4 time points, for 15 *ani-1;pig-1* embryos with corrected DNA segregation defects. All time points are shown on plots, t-test and regression test are performed on subsamples for which velocities are calculated from disjoint time windows.

#### Cortical tension measurement

Analysis of cortical ablation experiments was performed by drawing an ellipse fitting the opening area (see Fig.5B) as previously described (Saha et al., 2016). Opening kinetics was analysed by measuring the increase of the ellipse minimum axis length following ablation. Cortical tension has been shown to be proportional to the velocity of cortex displacement which immediately follows ablation (Mayer et al., 2010). Here, the initial opening velocity was calculated by measuring the increase of the ellipse minimum axis between the first (0.5 sec) and third (1.5 sec) images taken after laser ablation.

#### Flow measurement

Particle Imaging Velocimetry (PIV) was used to measure cytoplasmic flows in strains expressing VIT-2::GFP to label yolk particles. PIV analysis was performed with the Matlab MPIV toolbox and the Minimum Quadric Differences algorithm. Images were analysed using interrogation windows of 32*32 pixels with 50% overlap. Measurements were performed in the furrow region during the last 100 seconds of furrow closure. Frames in which nucleus displacement may have influenced the calculation of cytoplasmic flow were excluded from the analysis. Intensities of the velocity vectors along the embryo antero-posterior axis were calculated; positive and negative values correspond to a flow towards the posterior and anterior pole of the embryos, respectively. The distribution of the measured vector intensities was then determined for each embryo and finally averaged among all observed embryos.

#### Modelling of the forces affecting cytokinetic furrow positioning

Assuming that cortical myosin results in a force *F* acting on the furrow and proportional to the difference of myosin concentration between anterior and posterior, ΔM, this force needs to be balanced with forces resisting furrow displacement (Fig.6B). Part of these forces are due to the resistance of the cortex itself, these can be assumed proportional to furrow velocity *v*, and can be modelled as −*αv* (Fig.6B). Another resistance to furrow motion is the drag force exerted by the cytoplasm, which, due to confinement, needs to flow through the cytokinetic ring in order to allow displacement of the furrow in the opposite direction (Fig.6B). This drag force will be equal to the dynamic pressure difference on each side of the furrow times the section area of the furrow, *R*^*2*^*−r*^*2*^, where *R* is the cell radius and *r* the cytokinetic ring radius. Assuming a Poiseuille flow, the dynamic pressure drop will scale as *vR*^*2*^*/r*^*4*^. The cytoplasmic drag force can thus be modelled as − *ß vR*^*2*^*/r*^*4*^ *(R*^*2*^ − *r*^*2*^). Finally, the equilibrium between the force exerted by myosin and the forces resisting furrow displacement can be expressed as F = *αv* + *ß vR*^*2*^*/r*^*4*^ *(R*^*2*^−*r*^*2*^). Hence, the furrow velocity will scale as v ∼ ΔM *r*^4^ /(*αr*^4^ + *ß R*^*2*^*(R*^*2*^−*r*^*2*^)), which we plot in Fig.6C and 6D. One aspect of the data not captured in this model is a drift velocity seemingly independent from both myosin and furrow opening, and which may be ascribed to another biological control of furrow positioning (e.g. local signalling of myosin recruitment and/or effect of blebs and cytoplasmic flow) adding up to the balance. When adding a constant drift velocity to the above model, adjusting the two parameters *α* and *β* allows to fit experimental datasets in the four curves shown in Fig. 6C and 6D. In those conditions, we find that *α*/*β*=0.03, indicating that cytoplasmic drag dominates other friction sources.

#### Modelling of the probability of bleb appearance

Following (Charras et al, 2008), we write the variations of free energy entailed by the nucleation of a bleb of height *δ* by the detachment from the cortex of a patch of membrane of radius *a*:

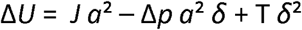

where *J* is the membrane-cytoskeleton energy of adhesion, Δ*p* the pressure jump across membrane, *T* the membrane tension; we have neglected bending energy.

In (Charras et al, 2008), the pressure jump is considered a uniform constant, since the cell is spherical and unconfined. In the present case, the confinement force adds up with the Laplace force and imposes a local curvature which is equal to the eggshell one (in locations where membrane is aposed to the eggshell) or larger than the eggshell one (at the furrow and at the poles when not in contact with eggshell). In these regions we have Δ*p* = *2σ/R*, where *σ* is the cortical tension and *R* the local characteristic radius of curvature. The lowest energy barrier to nucleate a bleb is at the saddle-point *δ*=JR/σ, a*=* (2TJ) ^½^R/*σ*, and is Δ*U* = *TJR/σ*. For a given cortical tension, the regions with smaller *R* (more curved) are thus more likely to nucleate blebs.

#### Statistical analysis

Statistical analysis and graphic representations were performed with Excel and RStudio softwares. Error bars represent standard deviation. Boxplots from Fig.5C and Fig.6A: central rectangles span the first to the third quartile of the data, middle lines correspond to median values, lower and higher whiskers to the minimum and maximum value, respectively. Fig.5C: two-tailed non parametric Wilcoxon tests were performed (data did not pass the Shapiro normality test). Fig.6A: a two-tailed unpaired Student’s t test was performed (data passed the Shapiro normality tests and sample variance was equal).

##### Key resources and experimental conditions tables

| Strain name | Genotype | Origin |
| --- | --- | --- |
| ANA058 | *pie-1p::GFP::TBA-2(ltIs25); pie-1p::GFP::H2B(oxIs279)* | Delattre's lab |
| ANA071 | *mex-5p::SPD-1::GFP(adIs1); pie-1p::mCherry::HIS-58(ltIs37); pie-1p::GFP::PH(PLC1delta1)(ltIs38)* | Delattre's lab |
| FL160 | *pie-1p::mCherry::HIS-58 (ltIs37); nmy-2::NMY-2::GFP(zuIs45)* | Cross OD56 with JJ1473 |
| FL178 | *pig-1(gm344)* | Remove *zdlIs5* from NG4370 |
| FL185 | *TBA-2::YFP(ddIs15) nmy-2(ne1490)* | Cross TH65 with WM180 |
| FL188 | *pig-1(gm344) pie-1p::mCherry::HIS-58(ltIs37); nmy-2::NMY-2::GFP(zuIs45)* | Cross FL178 with FL160 |
| FL191 | *TBA-2::YFP(ddIs15) nmy-2(ne1490); pig-1(gm344)* | Cross FL178 with FL185 |
| FL197 | *dhc-1(or195); pig-1(gm344) pie-1p::mCherry::HIS-58(ltIs37); nmy-2::NMY-2::GFP(zuls45)* | Cross EU828 with FL188 |
| FL201 | *emr-1p::EMR-1::mCherry(bqSi142); nmy-2::NMY-2::GFP(zuls45)* | Cross BN147 with FL188 |
| FL202 | *emr-1p::EMR-1::mCherry(bqSi142); pig-1(gm344); nmy-2::NMY-2::GFP(zuls45)* | Cross BN147 with FL188 |
| FL207 | *zyg-12(or577); pie-1p::mCherry::HIS-58(ltIs37); nmy-2::NMY-2::GFP(zuls45)* | Cross EU1590 with FL160 |
| FL208 | *zyg-12(or577); pig-1(gm344) pie-1p::mCherry::HIS-58(ltIs37); nmy-2::NMY-2::GFP(zuls45)* | Cross EU1590 with FL188 |
| FL209 | *spd-1(oj5); pie-1p::mCherry::HIS-58(ltIs37); nmy-2::NMY-2::GFP(zuls45)* | Cross FL160 with WH12 |
| FL212 | *TBA-2::YFP(ddIs15); pie-1p::mCherry::HIS-58(ltIs37)* | Cross TH65 with FL197 |
| FL213 | *nmy-2(ne1490); pig-1(gm344) pie-1p::mCherry::HIS-58(ltIs37)* | Cross FL188 with FL191 |
| FL214 | *pig-1(gm344) pie-1p::mCherry::HIS-58(ltIs37)* | Cross FL188 with FL191 |
| FL216 | *TBA-2::YFP(ddIs15); pig-1(gm344) pie-1p::mCherry::HIS-58(ltIs37)* | Cross TH65 with FL197 |
| FL217 | *TBA-2::YFP(ddIs15); zyg-12(or577); pie-1p::mCherry::HIS-58(ltIs37)* | Cross TH65 with FL208 |
| FL218 | *TBA-2::YFP(ddIs15); zyg-12(or577); pig-1(gm344) pie-1p::mCherry::HIS-58(ltIs37)* | Cross TH65 with FL208 |
| FL220 | *pig-1(gm344) pie-1p::mCherry::HIS-58(ltIs37); pie-1p::GFP::PH(PLC1delta1)(ltIs38)* | Cross TH155 with FL188 |
| FL221 | *nmy-2(ne1490); pie-1p::mCherry::HIS-58(ltIs37)* | Cross FL188 with WM180 |
| FL223 | *spd-1(oj5); pig-1(gm344) pie-1p::mCherry::HIS-58(ltIs37); nmy-2::NMY-2::GFP(zuls45)* | Cross FL188 with FL209 |
| FL225 | *nmy-2(ne1490); pig-1(gm344) pie-1p::mCherry::HIS-58(ltIs37); VIT-2::GFP(pwIs23)* | Cross FL213 with RT130 |
| FL227 | *pig-1(gm344) pie-1p::mCherry::HIS-58(ltIs37); VIT-2::GFP(pwIs23)* | Cross FL213 with RT130 |
| FL228 | *pie-1p::mCherry::HIS-58(ltIs37); VIT-2::GFP(pwIs23)* | Cross FL213 with RT130 |
| FL229 | *zyg-12(or577); pig-1(gm344) pie-1p::mCherry::HIS-58(ltIs37)* | Cross FL213 with FL218 |
| FL230 | *nmy-2(ne1490); zyg-12(or577); pig-1(gm344) pie-1p::mCherry::HIS-58(ltIs37)* | Cross FL213 with FL218 |
| FL233 | *nmy-2(ne1490); pig-1(gm344) pie-1p::mCherry::HIS-58(ltIs37); pie-1p::GFP::PH(PLC1delta1)(ltIs38)* | Cross FL221 with FL220 |
| FL235 | *pie-1p::GFP::TBA-2(ltIs25); pie-1p::GFP::H2B(oxIs279); nmy-2::NMY-2::GFP(zuls45)* | Cross ANA0058 with FL160 |
| FL256 | *mex-5p::SPD-1::GFP(adels1); pig-1(gm344) pie-1p::mCherry::HIS-58(ltIs37); pie-1p::GFP::PH(PLC1delta1)(ltIs38)* | Cross ANA0071 with FL220 |
| OD56 | *unc-119(od3); pie-1p::mCherry::HIS-58(ltIs37)* | Oegema's lab |

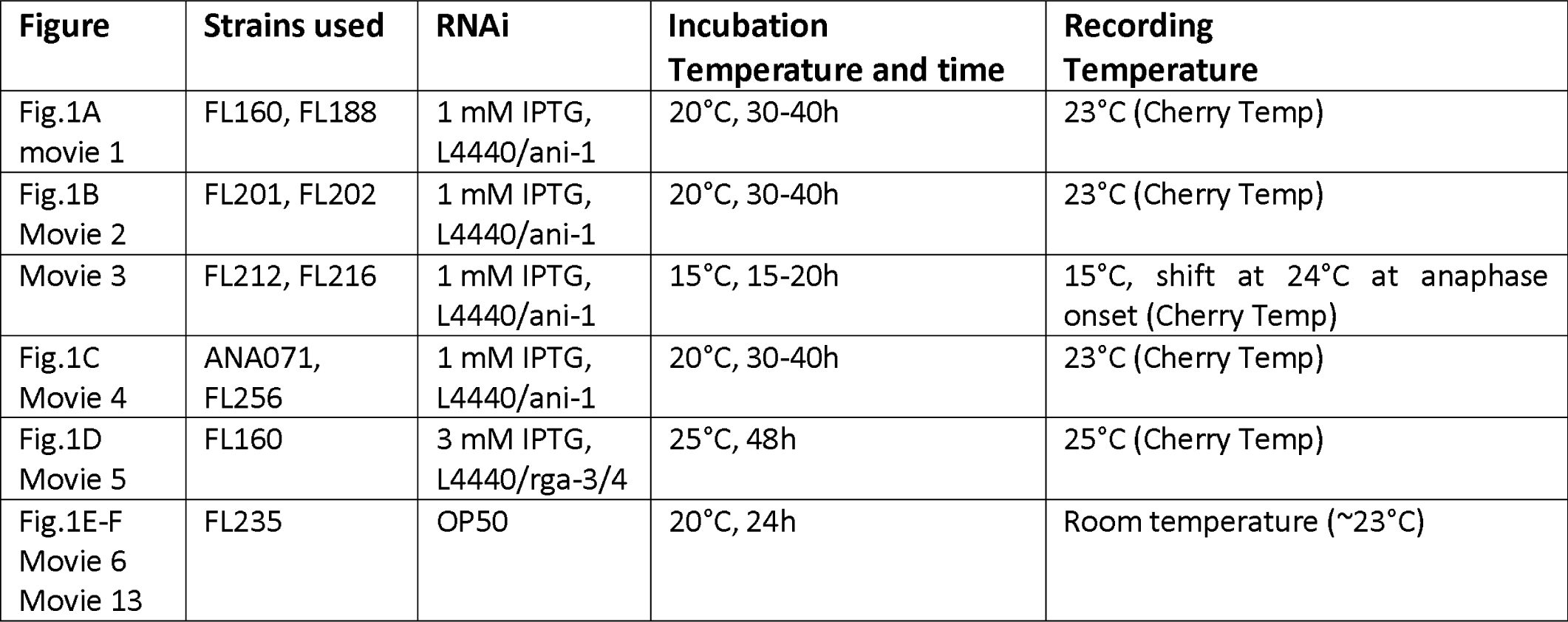
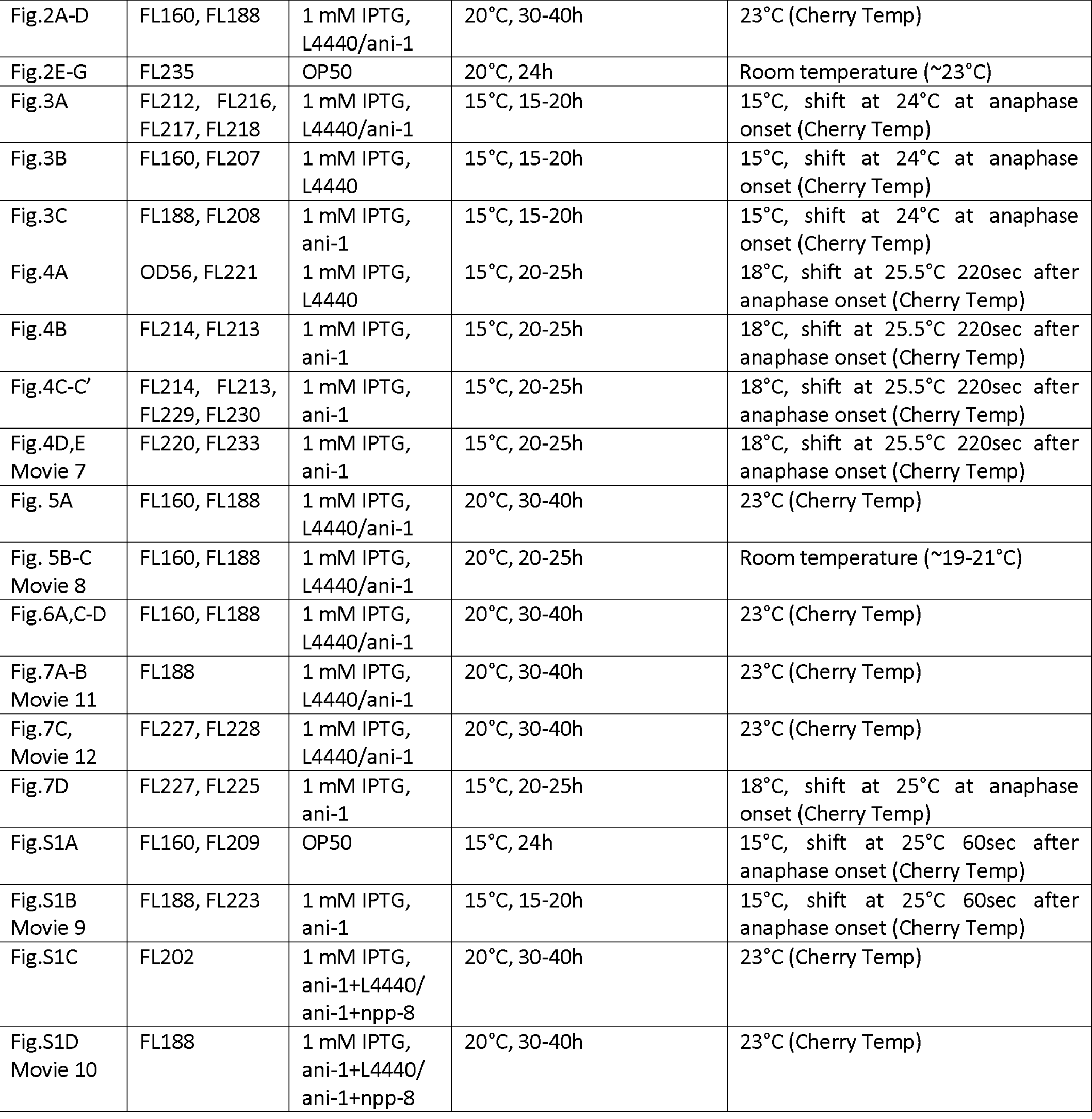

## Supplemental information

**Figure S1 (related to Figure 7):**
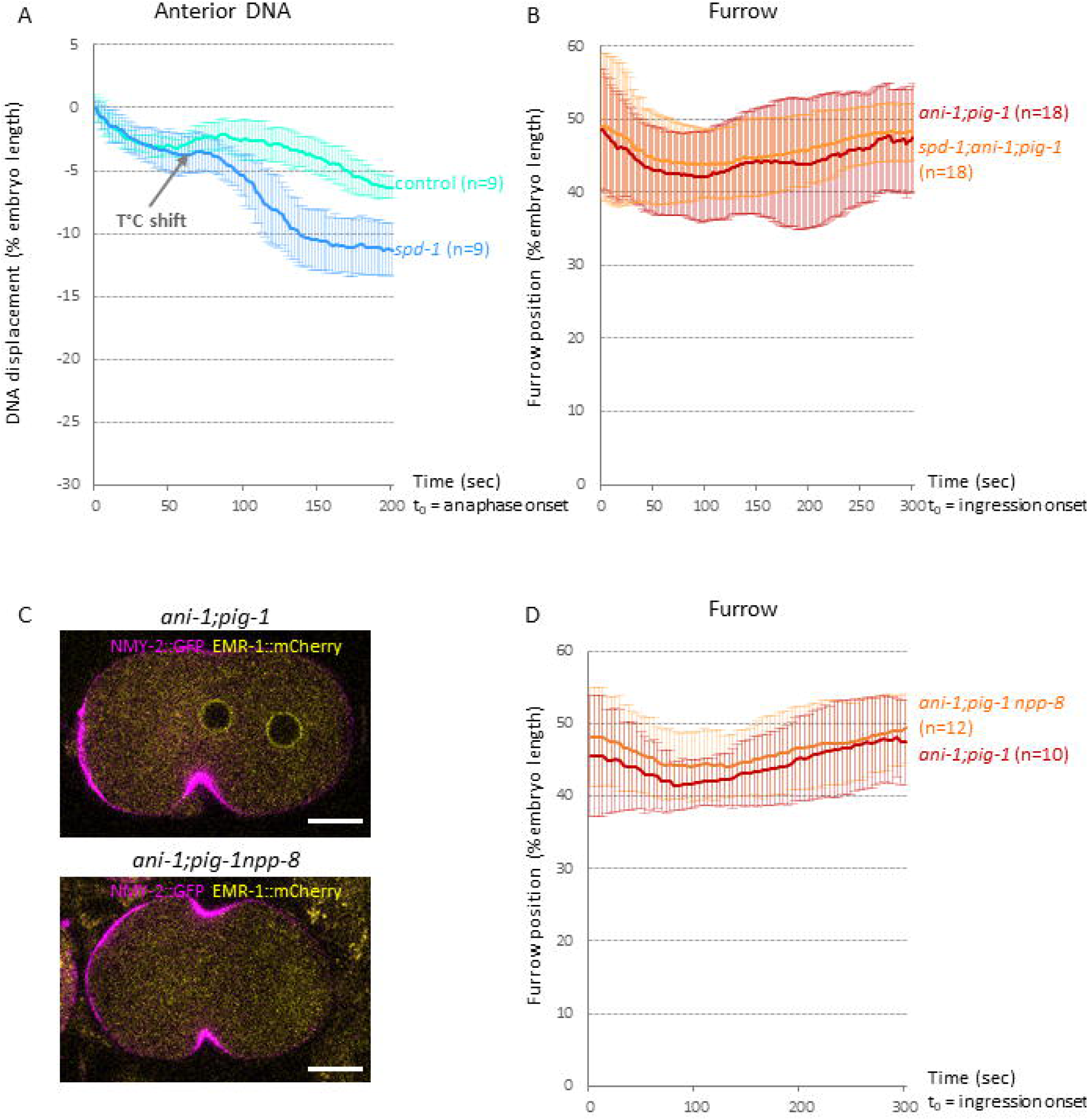
Central spindle and nuclear envelop formation are not required for furrow displacement. **A.** Average displacement of the anterior nucleus in control and *spd-1(oj5)* embryos expressing NMY-2::GFP and mCherry::HIS. In *spd-1(oj5)* embryos, anterior nucleus displacement is accelerated upon temperature shift, indicating efficient central spindle inactivation. Positive and negative values correspond to a displacement towards the posterior and anterior pole of the embryos, respectively; distances are expressed in percentage of embryo length. t_0_ = anaphase onset. **B.** Average furrow tip position along the antero-posterior axis in *ani-1(RNAi);pig-1(gm344)* and *spd-1(oj5);ani-1(RNAi);pig-1(gm344)* embryos expressing NMY-2::GFP and mCherry::HIS. Only embryos with corrected DNA segregation defects were analysed. All displayed cortical blebbing on the anterior side of the furrow. Rescue efficiency was similar for *ani-1(RNAi);pig-1(gm344)* and *spd-1(oj5);ani-1(RNAi);pig-1(gm344)* embryos (75% and 73% rescue, respectively). **C.** *ani-1(RNAi);pig-1(gm344)* and *ani-1(RNAi);pig-1(gm344)npp-8(RNAi)* embryos expressing NMY-2::GFP (magenta) and EMR-1::mCherry (yellow). No nuclear envelope reformation is observed when *npp-8* is depleted. Embryos are oriented with the anterior to the left. Scale bar: 10 μm. **D.** Average furrow tip position along the antero-posterior axis in *ani-1(RNAi);pig-1(gm344)* and *ani-1(RNAi);pig-1(gm344)npp-8(RNAi)* embryos expressing NMY-2::GFP and mCherry::HIS. Only embryos with corrected DNA segregation defects were analysed. *ani-1(RNAi);pig-1(gm344)* and *ani-1(RNAi);pig-1(gm344)npp-8(RNAi)* embryos present similar frequencies of cortical blebbing on the anterior side of the furrow (90% and 83% of rescued embryos, respectively) and similar rescue efficiency (91% and 100% rescue, respectively). 100 and 0 correspond to the posterior and anterior pole of the embryos, respectively. t_0_ = furrow ingression onset. In A, B, D: error bars correspond to standard deviation.

**Movie 1 (related to Fig.1A): Correction of DNA segregation defects resulting from furrow mispositioning in *ani-1:pia-l* embryos.** Control (left) and *ani-1(RNAi);pig-1(gm344)* (right) embryos expressing NMY-2::GFP (magenta) and mCherry::HIS (yellow). Increased myosin levels at the anterior cortex of *ani-1(RNAi);pig-1(gm344)* embryos lead to furrow mispositioning and DNA segregation defects which are corrected late during mitosis. In control embryos the anterior nucleus (yellow asterix) is slightly displaced during cytokinesis and furrow position remains stable (arrow). Correction in *ani-1(RNAi);pig-1(gm344)* embryos involves the strong displacement of the anterior nucleus (yellow asterix) to the anterior and the displacement of the furrow (arrow) to the posterior. Images were recorded at 4-s intervals and movies are played at 10 frames per second. t_0_ = anaphase onset. Embryos are oriented with the anterior to the left. Scale bar: 10 μm.

**Movie 2 (related to Fig.1B): Correction of DNA segregation defects in *ani-1;pig-1* embryos occurs after nuclear envelope reformation.** Control (left) and *ani-1(RNAi);pig-1(gm344)* (right) embryos expressing NMY-2::GFP (magenta) and EMR-l::mCherry (yellow). Nuclear envelope reassembles before the correction of DNA segregation defects. Images were recorded at 5-s intervals and movies are played at 10 frames per second. t_0_ = nuclear envelope breakdown. Embryos are oriented with the anterior to the left. Scale bar: 10 μm.

**Movie 3 (related to Fig.1C): Correction of DNA segregation defects in *ani-1;pia-l* embryos occurs after mitotic spindle disassembly.** Control (left) and *ani-1(RNAi);pig-1(gm344)* (right) embryos expressing a-tubulin::YFP (magenta in upper images, gray in lower images) and mCherry::HIS (yellow in upper images). The mitotic spindle starts disassembling before the correction of DNA segregation defects. Images were recorded at 5-s intervals and movies are played at 10 frames per second. t_0_ = anaphase onset. Embryos are oriented with the anterior to the left. Scale bar: 10 μm.

**Movie 4 (related to Fig.1C): Correction of DNA segregation defects in *ani-1:pia-l* embryos occurs after central spindle disassembly.** Control (left) and *ani-1(RNAi);pig-1(gm344)* (right) embryos expressing SPD-1::GFP (magenta, labels the central spindle, centrosomes and the nuclei after division), PH::GFP (magenta, membrane labelling) and mCherry::HIS (yellow). The central spindle starts disassembling before the correction of DNA segregation defects. Note that some central spindle remnants are displaced towards the anterior by the cytoplasmic flow. Images were recorded at 5-s intervals and movies are played at 10 frames per second. t_0_ = anaphase onset. Embryos are oriented with the anterior to the left. Scale bar: 10 μm.

**Movie 5 (related to Fig.1D): Correction of DNA segregation defects resulting from furrow mispositioning in *rga-3/4* embryos.** Control (left) and *rga-3/4(RNAi)* (right) embryos expressing NMY-2::GFP (magenta) and mCherry::HIS (yellow). Increased myosin levels at the anterior cortex of *rga-3/4(RNAi)* embryos lead to furrow mispositioning and DNA segregation defects which are corrected late during mitosis. Correction involves the opposite displacement of the anterior nucleus (yellow asterix) and the furrow (arrow) towards the anterior and posterior pole of the embryo, respectively. Images were recorded at 1.9-s intervals and movies are played at 10 frames per second. t_0_ = anaphase onset. Embryos are oriented with the anterior to the left. Scale bar: 10 μm.

**Movie 6 (related to Fig.1E-F): Correction of DNA segregation defects following centrosome ablations.** Ablation of anterior (left) or posterior (right) centrosome (white circle) in embryos expressing NMY-2::GFP (green, cortical and furrow signals), GFP:: α-tubulin (green, labels the centrosome and to a lesser extent microtubules) and GFP::HIS (green, labels the DNA). Centrosome ablation leads to spindle and/or furrow mispositioning. Correction of the resulting DNA segregation defects involves both nucleus (asterix) and furrow (arrow) displacement. In the case of anterior centrosome ablation (left), the anterior nucleus moves to the anterior and the furrow to the posterior of the embryo. Reversely, in the case of posterior centrosome ablation (right), the posterior nucleus moves to the posterior and the furrow to the anterior of the embryo. Images were recorded at 2-s intervals and movies are played at 10 frames per second. t_0_ = anaphase onset. Embryos are oriented with the anterior to the left. Scale bar: 10 μm.

**Movie 7 (related to Fig.4D): Myosin activity controls furrow displacement.** *ani-1 (RNAi);pig-1(gm344)* (left) and *nmy-2(ne1490);ani-1(RNAi);pig-1(gm344)* (right) embryos expressing GFP::PH (magenta) and mCherry::HIS (yellow, anterior nucleus also marked with asterix). During correction, the furrow is displaced towards the posterior in *ani-1(RNAi);pig-1(gm344)* embryos but not in *nmy-2(ne1490);ani-1(RNAi);pig-1(gm344)* embryos. In *ani-1(RNAi);pig-1(gm344)* embryos, furrow displacement is associated with cortical blebbing on the anterior side of the furrow (white arrowheads). t_0_ = anaphase onset. Images were recorded at 2-s intervals and movies are played at 10 frames per second. Embryos are oriented with the anterior to the left. Scale bar: 10 μm.

**Movie 8 (related to Fig.5B): Laser-induced cortical ablation.** Ablation of the anterior cortex of a *ani-1(RNAi);pig-1(gm344)* embryo was performed during anaphase. The embryo expresses NMY-2::GFP (green) and mCherry::HIS (not shown). White bar indicates the area targeted for laser ablation. Movie starts 3 sec before laser ablation. Images were recorded at 0.5-s intervals and movie is played at 10 frames per second. Scale bar: 5 μm.

**Movie 9 (related to Fig.S1B): SPD-1 inactivation does not prevent furrow displacement.** DIC recording of a *spd-1(oj5);ani-1(RNAi);pig-1(gm344)* embryo expressing NMY-2::GFP and mCherry::HIS (not shown). SPD-1 inactivation does not prevent cortical blebbing (arrowheads) or furrow displacement (arrow). to = furrow ingression onset. Images were recorded at 2-s intervals and movie is played at 10 frames per second. Embryo is oriented with the anterior to the left. Scale bar: 10 μm.

**Movie 10 (related to Fig.S1D): NPP-8 depletion does not prevent furrow displacement.** *ani-1(RNAi);pig-1(gm344)npp-8(RNAi)* embryo expressing NMY-2::GFP (magenta) and mCherry::HIS (yellow). NPP-8 depletion does not prevent cortical blebbing (arrowheads) or furrow displacement (arrow). t_0_ = furrow ingression onset. Images were recorded at 4-s intervals and movie is played at 10 frames per second. Embryo is oriented with the anterior to the left. Scale bar: 10 μm.

**Movie 11 (related to Fig.7A): *ani-1;pig-1* embryos display strong cortical blebbing.** DIC (left) and GFP (right) images of a *ani-1(RNAi);pig-1(gm344)* embryo expressing NMY-2::GFP (gray, right images) and mCherry::HIS (not shown). Cortical blebs first form both at the posterior and anterior cortical poles (arrows) and then on the anterior side of the cytokinetic furrow (arrowheads). Images were recorded at 4-s intervals and movies are played at 10 frames per second. t_0_ = anaphase onset. Embryo is oriented with the anterior to the left. Scale bar: 10 μm.

**Movie 12 (related to Fig.7C): Correction of DNA segregation defects in *ani-1;pig-1* embryos is associated with strong cytoplasmic flow.** Control (left) and *ani-1(RNAi);pig-1(gm344)* (right) embryos expressing VIT-2::GFP (green). A strong flow of cytoplasmic particles directed towards the anterior pole is observed during the correction of DNA segregation defects in *ani-1(RNAi);pig-1(gm344)* embryos. t_0_ = cytokinetic ring closure. Images were recorded at 1-s intervals and movies are played at 10 frames per second. Embryos are oriented with the anterior to the left. Scale bar: 10 μm.

**Movie 13 (related to Fig.7C): Correction of DNA segregation defects following centrosome ablations is associated with cytoplasmic flow.** Ablation of anterior (left) or posterior (right) centrosome (white circle) in embryos expressing NMY-2::GFP (green, cortical and furrow signals), GFP:: *α*-tubulin (green, labels the centrosome and to a lesser extent microtubules) and GFP::HIS (green, labels the DNA). Cytoplasmic particle movements can be observed on DIC images. In the case of anterior centrosome ablation (left), the anterior nucleus (asterix) and cytoplasmic particles (arrowhead) move to the anterior while the furrow (arrow) moves to the posterior of the embryo (4/4 embryos). In the case of posterior centrosome ablation (right), the posterior nucleus (asterix) and cytoplasmic particles (arrowhead) move to the posterior while the furrow (arrow) moves to the anterior of the embryo (5/6 embryos). Images were recorded at 2-s intervals and movies are played at 10 frames per second. t_0_ = anaphase onset. Embryos are oriented with the anterior to the left. Scale bar: 10 μm.

